# AGILE Platform: A Deep Learning-Powered Approach to Accelerate LNP Development for mRNA Delivery

**DOI:** 10.1101/2023.06.01.543345

**Authors:** Yue Xu, Shihao Ma, Haotian Cui, Jingan Chen, Shufen Xu, Kevin Wang, Andrew Varley, Rick Xing Ze Lu, Bo Wang, Bowen Li

**Author notes:** These authors contributed equally (Yue Xu, Shihao Ma, Haotian Cui). Communication can be sent to (Bo Wang, Bowen Li).

## Abstract

Ionizable lipid nanoparticles (LNPs) have seen widespread use in mRNA delivery for clinical applications, notably in SARS-CoV-2 mRNA vaccines. Despite their successful use, expansion of mRNA therapies beyond COVID-19 is impeded by the absence of LNPs tailored to different target cell types. The traditional process of LNP development remains labor-intensive and cost-inefficient, relying heavily on trial and error. In this study, we present the **A**I-**G**uided **I**onizable **L**ipid **E**ngineering (AGILE) platform, a synergistic combination of deep learning and combinatorial chemistry. AGILE streamlines the iterative development of ionizable lipids, crucial components for LNP-mediated mRNA delivery. This approach brings forth three significant features: efficient design and synthesis of combinatorial lipid libraries, comprehensive in silico lipid screening employing deep neural networks, and adaptability to diverse cell lines. Using AGILE, we were able to rapidly design, synthesize, and evaluate new ionizable lipids for mRNA delivery in muscle and immune cells, selecting from a library of over 10,000 candidates. Importantly, AGILE has revealed cell-specific preferences for ionizable lipids, indicating the need for different tail lengths and head groups for optimal delivery to varying cell types. These results underscore the potential of AGILE in expediting the development of customized LNPs. This could significantly contribute to addressing the complex needs of mRNA delivery in clinical practice, thereby broadening the scope and efficacy of mRNA therapies.

**One Sentence Summary:** AI and combinatorial chemistry expedite ionizable lipid creation for mRNA delivery.

## Introduction

Messenger RNA (mRNA) has emerged as a versatile tool with wide-ranging biomedical applications, ranging from vaccines and protein replacement therapy to cell engineering and gene editing ^1, 2^. This versatility has fueled widespread interest in exploiting mRNA to tackle an array of diseases ^3, 4^. However, the inherently unstable nature of mRNA and its susceptibility to nuclease degradation necessitates an effective delivery system, a role typically fulfilled by ionizable lipid nanoparticles (LNPs) ^5^. Both Comirnaty and Spikevax, two SARS-CoV-2 vaccines approved amidst the COVID-19 pandemic, are grounded on LNP-based mRNA delivery ^6, 7^. Moreover, LNP technology helped the first siRNA drug (Onpattro) obtain U.S. FDA approval in 2018 ^8–10^. The classical LNP formulation comprises four compositions: ionizable lipids, cholesterol, helper lipids, and PEGylated lipids. Notably, each of the three FDA-approved RNA LNPs has a distinct ionizable lipid design, highlighting the pivotal role of ionizable lipids in LNP technology. Their primary functions include packaging mRNA into LNPs and facilitating its entry into the cytoplasm of target cells for ribosomal binding and subsequent protein expression ^11–14^. An ionizable lipid generally consists of an ionizable amine head group and two lipid tails. This structure enables protonation at acidic pH, thereby adopting a cationic character during the LNP formulation process, facilitating the encapsulation of anionic RNA molecules. At physiological pH, the ionizable lipid remains neutrally charged, thereby circumventing potential toxicity associated with non-ionizable cationic lipids. Once the LNP encapsulating mRNA is endocytosed, ionizable lipids undergo protonation again in the acidic endosomal environment, disrupting the inner phospholipid membrane of endosomes and promoting the release of mRNA into the cytoplasm of target cells. As the COVID-19 pandemic recedes, the spectrum of mRNA applications continues to broaden beyond vaccination, thus emphasizing the necessity for a diverse array of ionizable lipids proficient in mRNA delivery to a variety of target cells and tissues.

Although previous research has provided some insight into the rational design of ionizable lipids to improve the mRNA delivery performance of LNPs, the approach often covers limited structural space, potentially overlooking some promising lipid designs. Combinatorial chemistry, employing multi-component reactions, has recently been used to enable high-throughput synthesis (HTS) of extensive and chemically diverse lipid libraries. For example, a Ugi-based three-component reaction (3-CR) could enable the swift synthesis of a combinatorial library, comprising 1,080 ionizable lipids, ultimately leading to the identification of a STING-activating ionizable lipid conducive to mRNA vaccine delivery ^15^. More recently, another 3-CR system based on the Michael addition was used to generate a library of over 700 ionizable lipids, resulting in the discovery of a potent lipid uniquely suited for efficient mRNA delivery to the lung epithelium ^16^. While the 3-CR combinatorial chemistry has been showcased to facilitate the synthesis of new ionizable lipids, constructing and testing a more extensive lipid library, running into hundreds of thousands of compounds, for mRNA transfection in different cell targets remains a formidable, time-consuming, and costly task ^17^. This challenge consequently restricts efforts to design and test more diverse and innovative structures. New strategies are essential to hasten the discovery and optimization of ionizable lipids for achieving desirable mRNA transfection in specific target cells.

Deep learning, a subset of artificial intelligence (AI), poses a promising resolution to the challenge of exploring molecular search spaces ^18–20^. With ample high-quality training data, these techniques can effectively extract insights from observed molecules, capitalizing on underlying chemical structures and properties, and extrapolating to a broader array of unobserved molecules. Indeed, the rise of deep learning is reshaping chemical compound discovery, transforming this process from a trial-and-error practice to an intelligent, data-driven strategy ^21–27^. In this study, we pioneered utilizing cutting-edge deep learning methodologies to accelerate the development of ionizable lipids for mRNA delivery, culminating in the **<u>A</u>**I-**<u>G</u>**uided **<u>I</u>**onizable **<u>L</u>**ipid **<u>E</u>**ngineering (AGILE) platform. This platform not only dramatically expands the molecular space of lipid structures by several magnitudes, but also significantly truncates the timeline for new ionizable lipid development, reducing it from potential months or even years to weeks. Essentially, AGILE employs a pre-trained deep-learning neural network that assimilates structural knowledge from millions of small-molecule components. The model utilizes vast amounts of unlabeled data from a combinatorial lipid library, employing a self-supervised approach to learn differentiable lipid representations. Following the fine-tuning on wet-lab data collected after HTS, AGILE can identify promising lipids for high mRNA transfection potency in specific cells from a significantly larger combinatorial library with enhanced accuracy. Leveraging this workflow, we fine-tuned the deep learning model using transfection data from Hela cells, which subsequently led to the prediction of 15 top lipid structures from a pool of 12,000 lipid candidates. This process facilitates the identification of an ionizable lipid H9 that shows superior mRNA transfection potency compared to LNPs containing (D-Lin-MC3-DMA) ^2^, an FDA-approved ionizable lipid for RNA delivery, following intramuscular injection. Notably, the transfection effect of H9 LNPs is localized to the muscle, with significantly less off-target transfection in other tissues, such as the liver. Moreover, we showed that AGILE could be quickly repurposed to discover LNPs for other target cells, as demonstrated by identifying a new lipid, R6, optimized for mRNA delivery to macrophages. Experimental observations, such as the significance of non-biodegradable tail structures in macrophage transfection and the correlation between the carbon chain length and transfection potency, underscore AGILE’s potential to provide meaningful biological insights and tailor LNPs for individual cell types. AGILE’s ability to customize for different cell types suggests its potential to steer the formulation of new mRNA-LNPs, finely tailored to various clinical scenarios.

## Results

### Overview of the AGILE platform

By synergistically integrating deep learning methodologies with combinatorial lipid synthesis chemistry, AGILE is dedicated to streamlining the discovery process for new ionizable lipids, which are crucial to LNP-based mRNA delivery. Central to this platform is a suite of deep learning algorithms, collectively referred to as the AGILE model. This model, encompassing a graph encoder and a molecular descriptor encoder, adeptly captures the intrinsic characteristics of ionizable lipid molecular structures and their corresponding chemical attributes. The implementation of AGILE in this study unfolds over three key stages, as illustrated in Figure 1a: (1) the constitution of a virtual library and initial self-supervised model training, (2) the acquisition of empirical data from an experimental library, enhancing the precision of the pre-trained model through supervised fine-tuning, and (3) the execution of *in silico* analysis on ionizable lipids in a candidate library, leveraging the refined deep learning algorithms (Methods 1.1 for additional details). As a multifaceted tool, AGILE generates predictions on the mRNA transfection capacity of ionizable lipids in LNP formulations and significantly facilitates the design of LNP for specific target cells.

**Figure 1.**
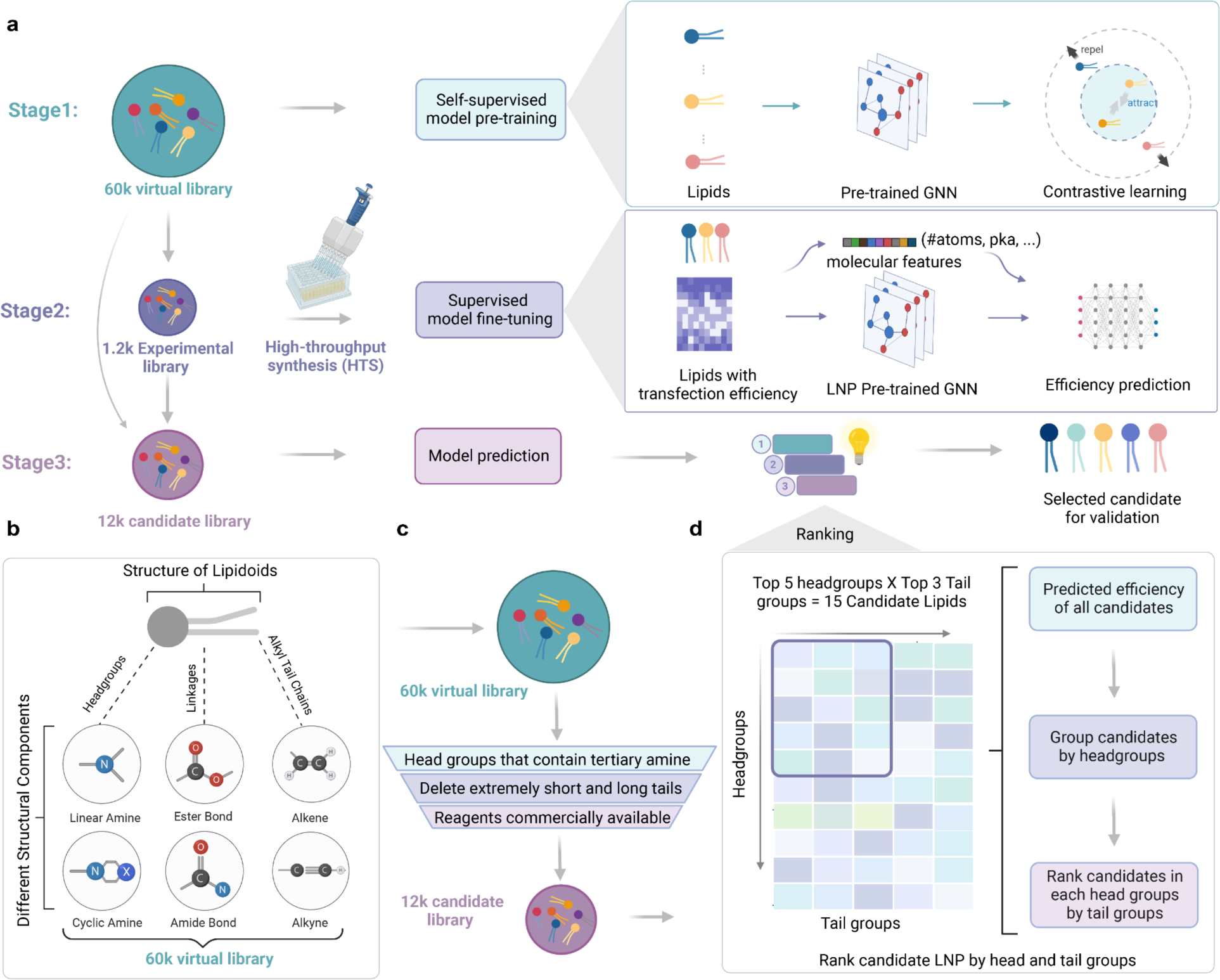
Overview of the platform design pipeline. (a) Illustration of the 3-stage workflow of the platform. Stage 1: Construction of a virtual library and self-supervised pre-training of the model. Stage 2: Synthesis of an experimental library for the fine-tuning of the model in a supervised manner. Stage 3: Deployment of the fine-tuned model for predictive analysis on a candidate library, followed by ranking for final candidate selection. (b) Depiction of virtual library design through the application of Ugi combinatorial chemistry. (c) Schematic representation of the rational selection process for lipid candidates, with 3 listed filtering criteria. (d) A comprehensive breakdown of the ranking procedure and the selection methodology for final candidates.

Stage 1 aims to develop a graph encoder proficient in differentiating and depicting distinct lipids through pre-training on a vast collection of unlabeled lipid molecules (Methods 1.3 and 1.4). This process begins with the construction of a graph encoder utilizing Graph Neural Networks (GNN), primed with parameters from the MolCLR model, which has undergone pre-training on a repertoire of over 10 million small molecules. This “warm-starting” strategy, embedding general knowledge of small molecular structures into our algorithm, fortifies the accuracy of AGILE in subsequent stages (Supplementary Fig. S1). The graph encoder subsequently underwent continuous pre-training on a virtual library of 60,000 chemically diverse lipids through contrastive learning ^28^, enabling the differentiation of atoms and bonds in each molecule, and thus capturing the disparities amongst various lipid structures (see Methods 1.4). This virtual library, composed of lipids with diverse amine head groups and two unique alkyl chains (Fig. 1b), is designed based on 3-CR chemistry principles, thus amenable to high-throughput combinatorial synthesis ^29^. Overall, the pre-training in Stage 1 equips the graph encoder with a comprehensive understanding of lipid structures, thereby enhancing subsequent steps (Supplementary Fig. S1). Stage 2 seeks to further train the AGILE model with mRNA transfection potency data from a pool of ionizable lipids. To this end, we synthesized 1200 ionizable lipids by 3-CR and assessed their transfection potency in a target cell line, from which the data was leveraged to fine-tune the AGILE model in a supervised manner (Methods 1.5). To enhance the generalizability and precision, we added a molecular descriptor encoder that takes molecular descriptors computed by Mordred as the input^30^ (Methods 1.3). The output of the molecular descriptor encoder was utilized to update the representation of lipid structures by the pre-trained graph encoder. As such, the AGILE model has been trained to minimize the difference between the predicted result and the ground truth from wet-lab experiments during the fine-tuning process. Prior to the *in silico* screening in Stage 3, we assembled a candidate library containing 12,000 lipid structures by rationally selecting structures from the virtual library in Stage 1 (Fig. 1c) following three rules (Methods 1.1): (1) Removal of non-ionizable cationic lipids due to the potential risk of toxicity ^31^; (2) Removal of lipids with too short (<C10) or too long (> C18) alkyl chains based on empirical experience ^15^; and (3) Removal of lipids requiring unavailable reagents for synthesis. The fine-tuned AGILE model was then utilized to predict the mRNA transfection potency of lipids in the candidate library, followed by a head and tail-wise ranking methodology to increase the structural diversity of the top-ranked candidates (Fig. 1d, Methods 1.6). Based on the information afforded by AGILE, the top-ranked ionizable lipid structures were selectively synthesized in the wet lab and formulated into LNPs for validating their ability to efficiently deliver mRNA to a specific target cell.

### Combinatorial Lipid library synthesis and screening for fine-tuning

Upon completing the pre-training of the entire virtual library in Stage 1, we tailored the model for transfection potency prediction through supervised fine-tuning. This stage involved training the model based on *in vitro* screening results, enabling the model to capture the potential transfection ability of molecules. To rapidly generate ionizable lipid libraries with high chemical diversity, we developed an automated high-throughput synthesis (HTS) platform based on the one-pot Ugi 3CR (Fig. 2a and Supplementary Fig. S2), which enabled the synthesis of a large batch (1,200) of ionizable lipids within 24 hours. The synthesized lipid library comprises 20 diverse head groups, 12 alkyl chains with biodegradable ester linkages, and 5 alkyl chains containing isocyanide function groups (Fig. 2b) ^32^. Using the HTS platform, we formulated LNPs via a liquid handling robot following a previously established classical four-composition formulation ratio ^33^.

**Figure 2.**
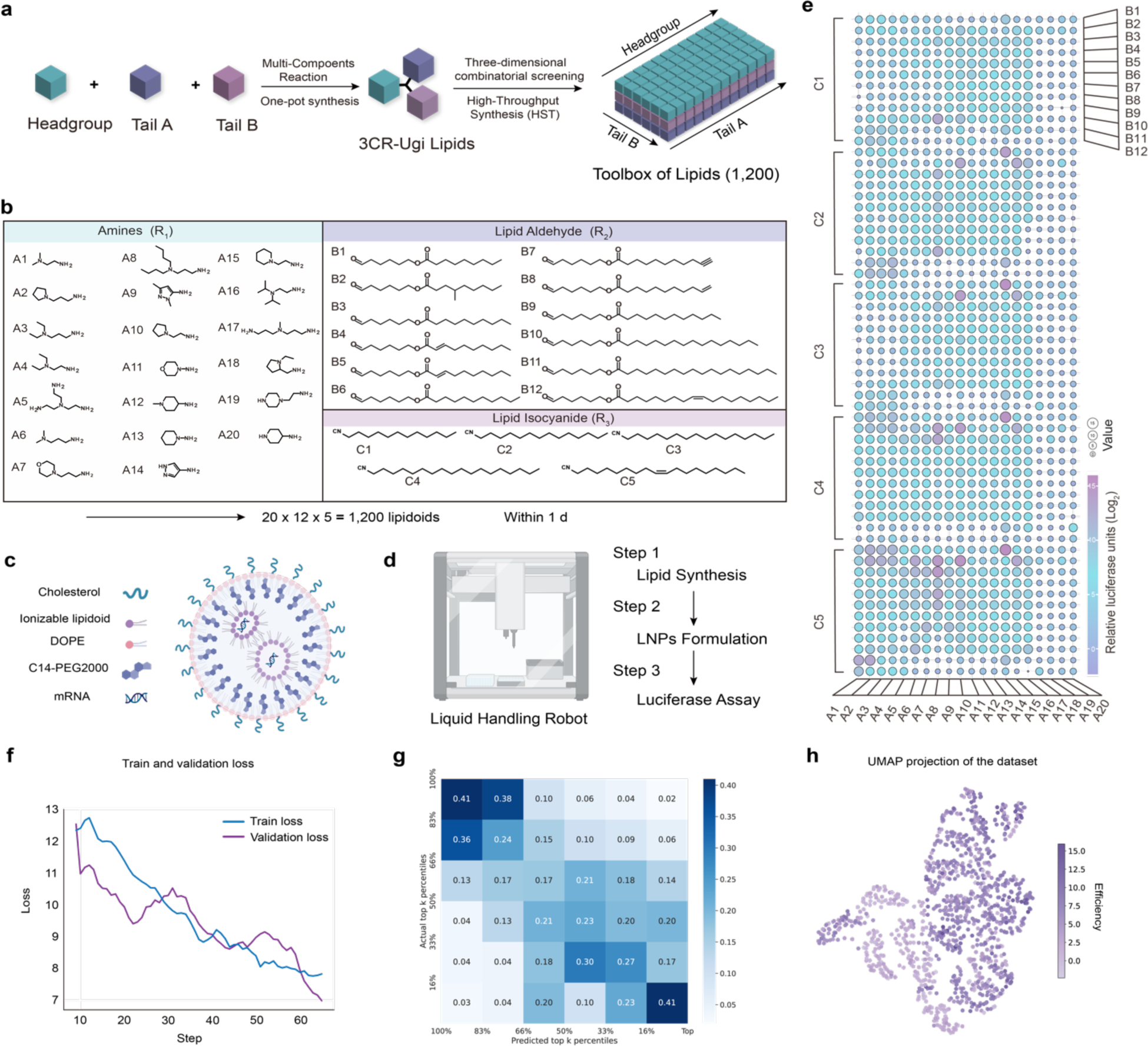
High throughput lipids synthesis and screening platform. (a) A schematic to illustrate the high-throughput synthesis method for lipids. (b) The combinatorial lipoids library consists of three components structure (amine head groups, aldehyde tails, and isocyanide tails). (c) A schematic diagram shows the LNPs components for mRNA encapsulating. (d) Lipid synthesis, LNPs formulation and luciferase assay based on liquid handling robot. (e) The data used for the fine-tuning are depicted in a balloon plot, which involved 1,200 LNPs for Fluc mRNA (mLuc) delivery and measuring the relative luciferase expression in Hela cells. (f) The loss value on the training set and validation set against fine-tuning steps. (g) The precision matrix computed on the experimental library of 1,200 lipids. The predicted and actual transfection potencies are divided into six equal percentiles. (h) UMAP plot of the experimental library, colored by the transfection potency.

The LNPs were subsequently synthesized for testing lead candidates by four classic formulations with the ionizable lipids, helper lipid (DOPE), cholesterol, and polyethylene glycol (PEG)-phospholipid conjugate (DMG-PEG2000) (Fig. 2c) ^34^. To evaluate the mRNA transfection potency in Hela cells, we measured firefly luciferase (Fluc) protein expression activity by encapsulating Fluc mRNA in the LNPs (Fig. 2d). Most of these 1200 lipids showed improved mRNA transfection potency in Hela cells compared to untreated cells (Fig. 2e). Hela cells are commonly utilized as an in vitro screening model for evaluating transfection potency through intramuscular injection. This is due to their reliable expression of the low-density lipoprotein receptor (LDLR), which plays a crucial role in the cellular uptake of lipid nanoparticles (LNPs) associated with lipoproteins in the bloodstream ^35^. The presence of LDLR in Hela cells allows for enhanced cellular uptake of LNPs, making them a valuable tool for assessing the effectiveness of intramuscular delivery methods. Previous studies emphasized the preference for muscle as the site of vaccination. This choice was based on the rich blood supply in muscle tissue, which enables the efficient processing of foreign antigens by immune cells, leading to a robust immune response ^36, 37^. Therefore, the strong correlation between transfection potency in Hela cells and in muscle tissues further establishes their utility in evaluating the effectiveness of intramuscular delivery methods ^39^. Meanwhile, the potencies vary significantly among test lipids, with relative luciferase units ranging from poor (Log_2_<5) to outstanding performance (Log_2_>10) (Supplementary Fig. S3). These variations can be readily used, in the fine-tuning stage, to supervise the model to learn the relation between molecule properties and its transfection potency (Methods 1.5). We used 80% of the data for the model training, 10% for selecting the best hyperparameters, and the last 10% for internal verification. We observed a constant decrease of loss value on both training and validate data (Fig. 2f) and thus used the model with the lowest validation loss.

To verify the quality of the predictions, we split the predicted and actual *in vitro* potency values into six equal percentiles. We visualize the precision matrix on all 1,200 lipids in Fig. 2g. Although the prediction task is extremely challenging, the model works particularly better for predicting the top and least performing lipids, which are arguably the most important and informative for selecting lipid candidates. For example, a predicted top-16% performing lipid will have a chance of 0.41 to be one of the actual top-16% performing lipids found *in vitro* (Fig. 2g). We also examined our predictions using UMAP embedding (Fig. 2h) ^38^. The UMAP algorithm assigns close LNPs presentations to adjacent points in a two-dimensional space, which are then colored based on their predicted transfection potency. The lipids gathered into regional structures with similar potency values on the resulting UMAP plot, which verifies that the learned representations capture the potential transfection ability of lipids.

### AGILE predicts and identifies the efficient lipid for muscle injection

With the fine-tuned model, we perform model prediction on the candidate library to screen potential lipids for muscle injection. We visualize our predictions using UMAP (Fig. 3a), and the resulting plot shows a clear separation between high and low predicted values, indicating the robustness of the model in differentiating efficacious and less efficacious ionizable lipids in a larger screening library. A closer look at the stratified distribution plots reveals that predicted potencies are clearly sorted by head group and tail combinations (Fig. 3b and Supplementary Fig. S4). Even among the top 5 performing head groups, A8 and A21 had higher predicted potencies than the others. While the tail combinations displayed less pronounced stratification of predicted transfection potencies compared to the head groups (Fig. 3c and Supplementary Fig. S4), the top tail combinations were still essential for candidate selection compared to the bottom tail combinations. The model appeared to favor unsaturated alkyl chains, a finding that was consistent with much of the literature that had been reported (Supplementary Fig. S5) ^39, 40^. Using our ranking system, which prioritizes structural diversity among lipids by considering head groups and tail combinations (Fig. 1d), we finalized a set of 15 lipid candidates (Supplementary Fig. S5).

**Figure 3.**
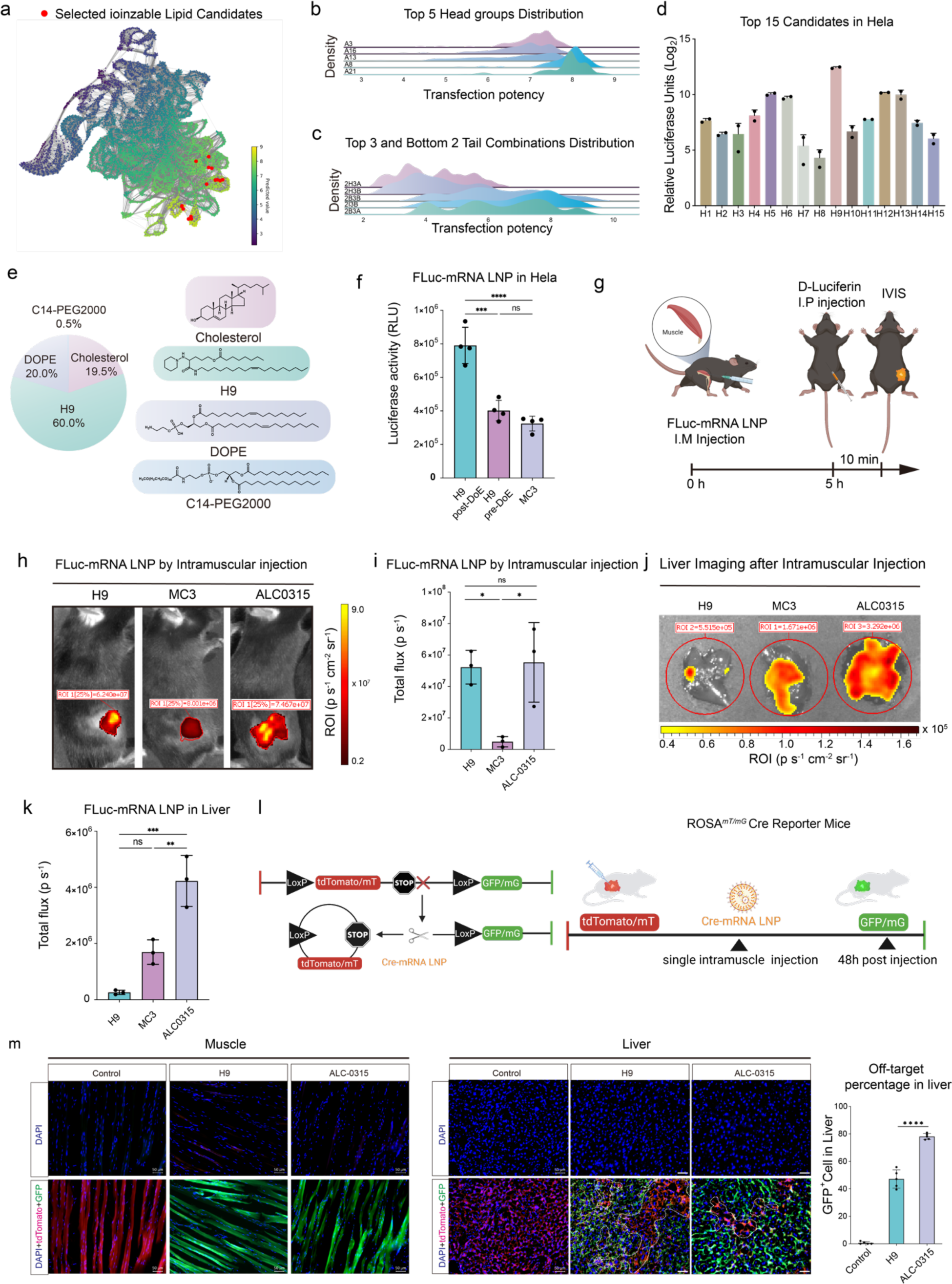
Model prediction and the validation of the gene editing potential with top-performing mRNA-LNPs. (a) The UMAP plot of the predicted molecule trans potencies. (b) Head group distribution and (c) tail combination distribution in Hela. (d) Validate 15 lipid candidates for Hela cell. (e) The top-performing formulation parameters used in the optimization of H9 LNPs in Hela. (f) Transfection of mFFL LNPs in Hela cells (n = 4 biologically independent experiments per group). (g) A schematic to illustrate the Intramuscular (IM) injection of mFluc-loaded LNPs into the mice and IVIS imaging. (h) LNPs formulated with FFL encoding mRNA were injected intramuscularly into mice (0.25mg mRNA/kg mouse). The top-performing lipids H9 with optimized formulation compare with the MC3 and ALC-0315 LNPs (n = 3 biologically independent mice per group, 0.5 mg kg⁻¹ mLuc per mouse). (i) Transfection of mFFL LNPs at the i.m. injection site in mice (n = 3 biologically independent mice per group). (j) IVIS imaging for liver after IM injection of mFluc-loaded LNPs. (k) Transfection of mFFL LNPs of liver in mice after IM injection (n = 3 biologically independent mice per group). (l) A schematic illustrating the Cre recombinase deletes STOP cassettes and activates the GFP mice reporter. (M) Representative confocal microscopy images and quantification of tdTomato and GFP expression in histological muscle and liver sections of mTmG mice post-injection of Cre-mRNA loaded LNPs by IM injection. Scale bar: 50µm. n = 5 sections from 3 mice. Error bars are S.D. Statistical significance was analyzed by the two-tailed Student’s t-test. ✱=p-value <0.05, ✱✱=p-value<0.01, ✱✱✱=p-value<0.005. Data are presented as mean±SD.

We rapidly synthesized the 15 lead candidates ranked by the model in the HTS system and evaluated them in Hela cells and found that all 15 lead candidates resulted in luciferase protein expression compared with the untreated group (Fig. 3d). To investigate their potential *in vivo*, we administered mice with Fluc mRNA encapsulated in LNPs by intramuscular injection. Among 15 different candidates, we observed a notably robust bioluminescence signal for H9 LNPs (Supplementary Fig. S6). After optimizing the LNPs formulation of H9 by using the design-of-experiment (DoE) (Fig. 3e, Supplementary Table. S1 and Fig. S7). After conducting a comparison with MC3 LNPs, we discovered that H9 LNPs had 2.3 times more mRNA transfection potency than MC3, which is a benchmark ionizable lipid currently used in the clinic (Fig. 3f) ^41^. Based on the positive outcome, we proceeded to use the H9 LNPs to assess mRNA transfection potency in mice through intramuscular injection (Fig. 3g). Our findings revealed that the transfection potency of the H9 LNPs in muscle site was 7.8 times stronger than that of the MC3, with no significant difference compared to the ALC-0315 (the ionizable lipid used in the SARS-CoV-2 vaccine, BNT162b2, from BioNTech and Pfizer) (Fig. 3h and i). It is worth noting that administering mRNA LNPs through intramuscular injection may cause an off-target effect, leading to the production of FLuc protein expression in the liver of mice ^42^. When compared to ALC-0315 LNPs, H9 LNPs were found to have lower off-target effects in the liver while maintaining similar transfection effectiveness in muscle tissue. (Fig. 3j and k). Inspired by these findings, we investigated the potential of H9 LNP for vaccination. To compare the delivering efficacy of H9 and ALC0315 LNPs, we administered cre-recombinase mRNA LNPs to mTmG reporter mouse models ^43^. These mice harbored gene mutations in the Gt(ROSA)26Sor locus, and upon cre-mRNA expression, the mT cassette was excised in the cre-expressing tissue, enabling the expression of the downstream membrane-targeted green fluorescent protein (GFP, mG) cassette (Fig. 3l). We observed comparable levels of GFP protein expression at the intramuscular injection site for H9 and ALC-0315 LNPs. However, ALC-0315 LNPs showed higher protein expression levels in liver tissue (Supplementary Fig. S8). Quantification of confocal images revealed that the H9 LNP exhibits 28% lower transfection potency in the liver compared to ALC-0315 LNPs (Fig. 3m). Notably, clinical studies have associated ALC-0315-based BNT162b2 mRNA vaccines with autoimmune hepatitis (AIH) following vaccination ^44^. Hence, it is anticipated that the H9 LNPs predicted by AGILE will alleviate the serious potential side effects of hepatitis with a lower off-targeting effect.

### Using AGILE to identify ionizable lipids for Macrophage mRNA delivery

It is known that conventional adeno-associated virus (AAV) vectors struggle to transfer innate immune cells, which highlights the importance of a non-viral mRNA delivery system ^45^. Although non-viral delivery vectors may avoid this disadvantage in immune cells, they still require effective mRNA transfection potency into the targeted immune cell type ^46^. In order to test AGILE’s ability to identify ionizable lipids that can efficiently transfect immune cells, we examined 1,200 lipids in RAW 264.7 cells (a macrophage cell line). This allowed us to create a dataset specifically for macrophages and fine-tune the screening process. The results revealed considerable differences in transfection potency between these two cell lines, with even the same batch of lipids showing totally disparate outcomes in Hela cells and macrophage cells (Supplementary Fig. S9). The study discovered that immune cells were less easily transfected by LNPs than Hela cells, demonstrating that immune cells pose a greater challenge for transfection. (Supplementary Fig. S10) ^47, 48^.

With the model fine-tuned on the macrophage-specific dataset, we once again performed model prediction and visualized the predicted transfection potencies for RAW 264.7 cells using UMAP (Fig. 4a). Contrasting with the UMAP of predicted potencies for Hela cells, the top-tier predicted LNPs are dispersed more widely throughout the space, potentially suggesting an increased complexity in predicting potencies for RAW 264.7 cells. Mirroring the pattern observed in Hela cells, the predicted potencies for RAW 264.7 cells exhibit evident stratification when categorized by head groups and tail combinations (Fig. 4b, c and Supplementary Fig. S4). The top 15 candidates were synthesized in the wet lab and subjected to an initial screen in RAW 264.7 cells, where 11 out of 15 showed improved transfection potency compared to MC3 (Fig. 4d and Supplementary Fig. S11). R6 was chosen as the best-performing lipid among the 15 candidates and subjected to formulation optimization using the design of experiments (DoE) (Fig.4e and Supplementary Fig. S12) ^49^. We then loaded LNPs with Fluc mRNA and evaluated luciferase protein expression in both RAW 264.7 and Hela cells to compare H9 and R6 LNPs performance in different cell lines. Interestingly, the results were quite different in RAW 264.7 cells, where R6 exhibited significantly higher transfection potency than H9 and MC3 (Fig. 4f). However, H9 demonstrated more than a 2-fold increase in transfection potency compared to R6 in Hela cells (Fig. 4g). These results demonstrated the necessity to develop LNPs specifically for individual cell types and tissues, rather than a one-size-fits-all approach for all targets. Based on the excellent performance of R6 LNPs in RAW 264.7, we tried to use R6 LNPs to deliver GFP mRNA to RAW 264.7. When compared to H9 and MC3 LNPs, R6 LNPs exhibited a 5-fold increase in transfection potency in RAW 264.7 as determined by flow cytometry (Fig. 4h and 4i). These results validated the success of AGILE in identifying a new ionizable lipid for efficient macrophage transfection, highlighting its potential to be utilized for the development of non-viral mRNA delivery vectors for immune cells.

**Figure 4.**
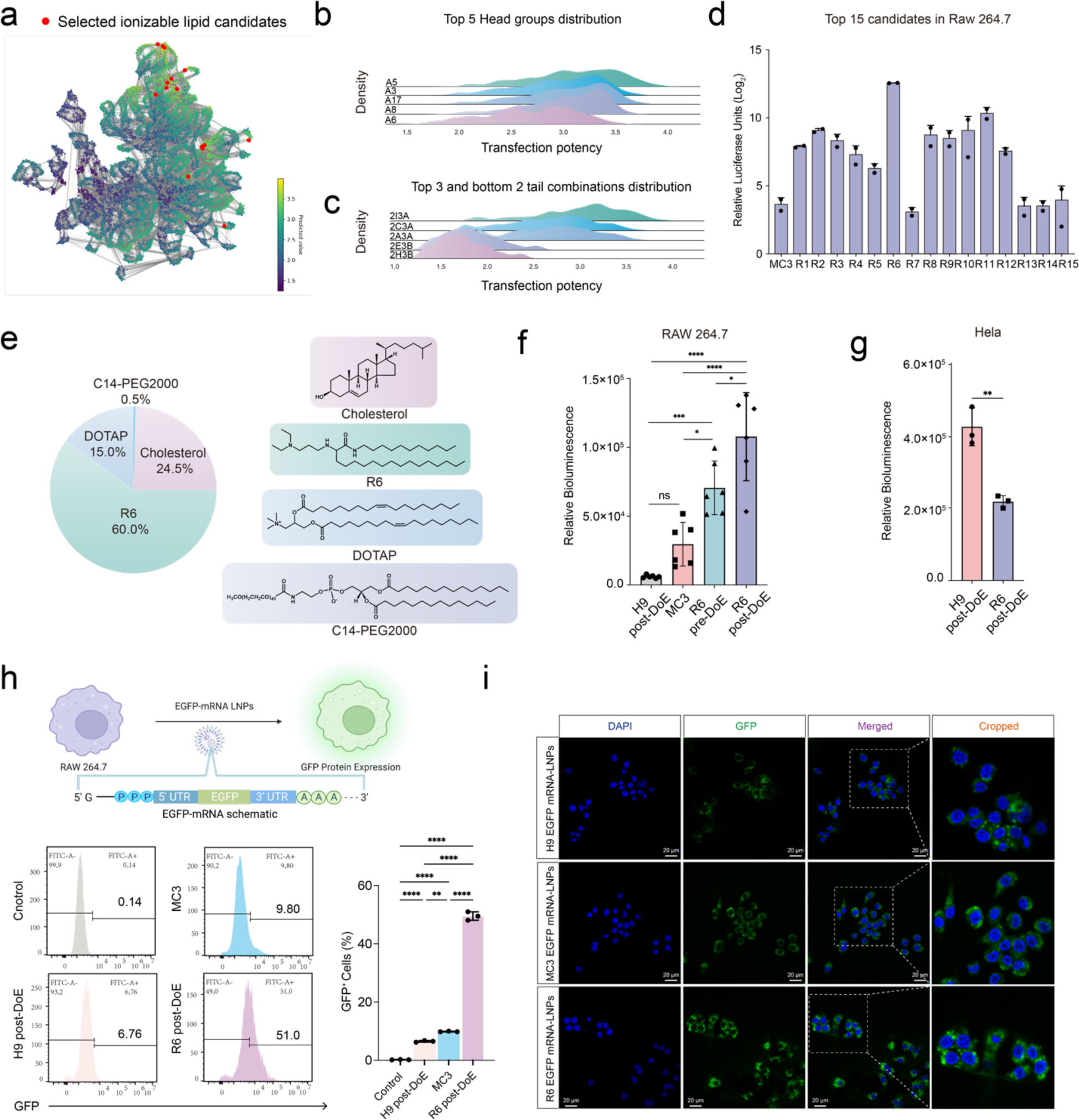
Accelerating screening of new lipids for EGFP-mRNA delivery in macrophage through the platform. (a) The UMAP plot of the predicted molecule trans potencies. (b) Top 5 Head groups distribution and (c) top 3 and bottom 2 tail combinations distribution in RAW 264.7. (d) Validate 15 lipid candidates for RAW 264.7 (n=2). (e) The top-performing formulation parameters used in the optimization of R6 LNPs in RAW 264.7. (f) Comparison of the Fluc-mRNA transfection potency of different LNPs in RAW 264.7 cells (n=6). (H9 LNPs, MC3 LNPs, R6 original screen formulation LNPs and optimized formulation LNPs). (g) Comparison of the efficacy of LNPs (H9 LNPs and R6 LNPs) in Hela cells (n=3). (h) Percentage of GFP positive cells on RAW 264.7 after treatment with MC3 LNPs, H9 LNPs and H6 LNPs. Quantitative analysis of flow cytometry data of RAW 264.7 cells (n=3). (i) Confocal images of RAW 264.7 cells transfected by GFP-mRNA LNPs. Green represents GFP, and blue represents the nucleus (DAPI). Statistical significance was analyzed by the two-tailed Student’s t-test. ✱=p-value <0.05, ✱✱=p-value<0.01, ✱✱✱=p-value<0.005. Data are presented as mean ± SD.

### Interpretation of the AGILE deep learning model

AGILE elucidates its models through two mechanisms: (1) identification of influential molecular descriptors using a gradient-based model interpretation method, and (2) discernment of critical features within selected lipids. We applied the gradient-based interpretation method to the 813 chosen molecular descriptors, assessing their contribution to the model’s prediction. As illustrated in Figures 5a and 5b, we have visualized the top 20 salient descriptors for both the Hela cell line and RAW 264.7. For the Hela cell line, VSA_EState3 and SssNH emerged as the most influential molecular characteristics for potency prediction. VSA_EState3, a descriptor quantifying the electronic and steric properties of a molecule’s surface area within a specific range^50^, along with SssNH, representing a tertiary amine, aligned with the expert understanding that head groups with tertiary amines are vital for lipid design. Subsequent analysis of essential features classified by head groups (Fig. 5k) pinpointed PEOE and Estate as the most critical descriptors for top-performing head groups (A13, A21), while SsNH2 (Sum of sNH2 E-states) and NsNH2 (Number of atoms of type sNH2) dominated in the least-performing groups (A5, A17) (Supplementary Fig. S4). Notably, these descriptors have strong associations with the amide bond in the structure, a critical connection within the 3CR Ugi Markush structure. This connection allows for various functional group attachments, influencing the lipid-like substances’ overall charge and their physicochemical properties within biological systems. Intriguingly, the model does not favor amide bond generation, potentially due to its impact on the overall physicochemical properties of lipids, such as pKa. In the context of RAW 264.7, SpDiam_Dzi and VR3_D are identified as the most influential descriptors (Fig. 5b). VSA_EState appears as the third most influential, implying its pivotal role in determining delivery potency to RAW 264.7, akin to Hela cells. Interestingly, head groups that underperformed in Hela (A5, A17) emerged as top performers in RAW 264.7, with SsNH2 and NsNH2 remaining the most critical features. In Hela cells, the cyclized head group outperformed the linear head group in transfection efficacy. However, the opposite trend was observed in RAW 264.7 cells. These observations underscore the necessity of designing LNPs with specific lipids tailored for distinct cellular targets.

**Figure 5.**
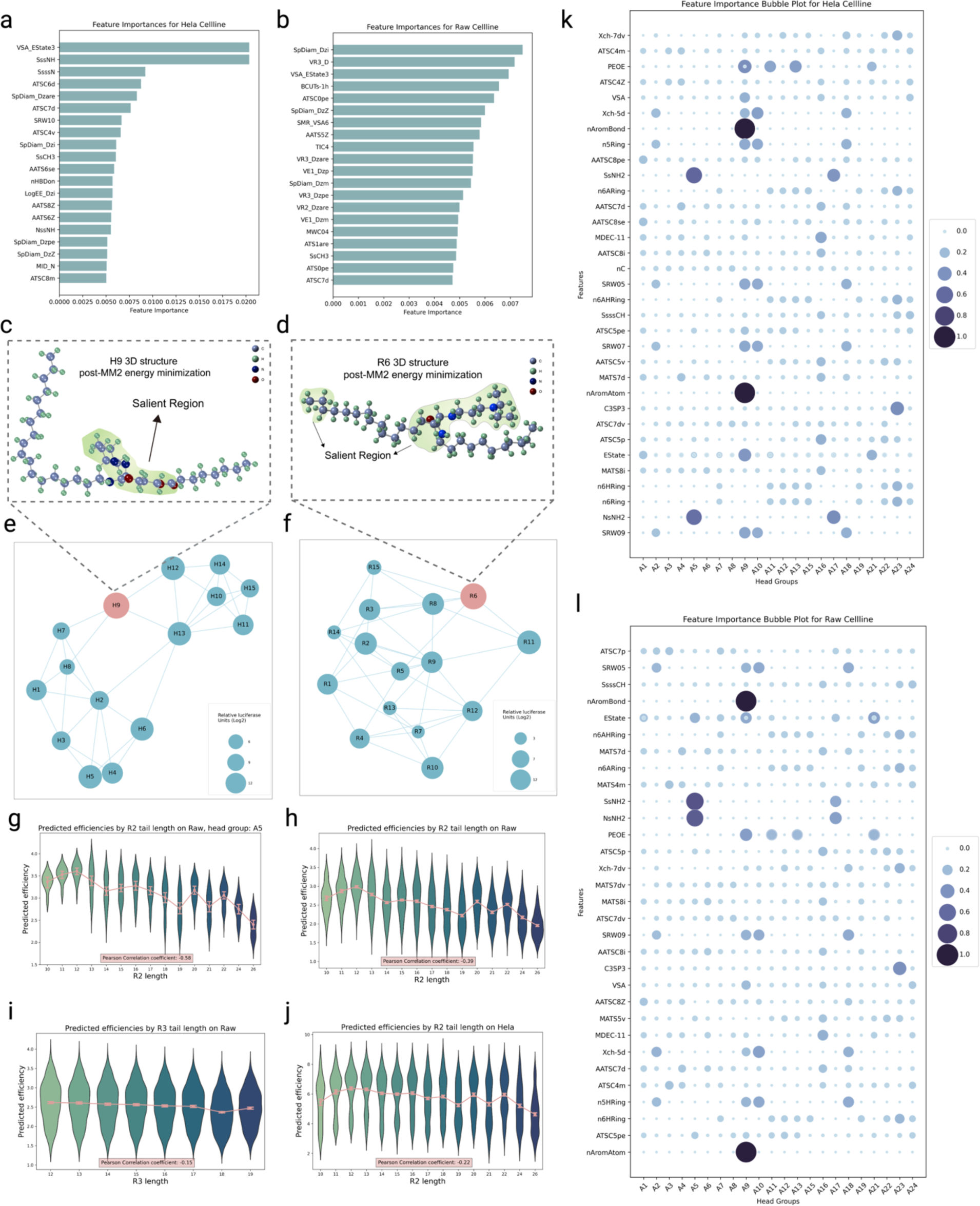
Model feature explanation and finding. (a, b) Top 20 most important molecular descriptors identified by this model fine-tuned for Hela and RAW 264.7 cell lines, respectively. (c, d) 3D visualization of H9 and R6 structures, respectively, with salient region highlighted. (e, f) Similarity networks for the 15 top lipid candidates in Hela and RAW 264.7 cell lines respectively, with each candidate linked to its four closest neighbors. (g) Violin plot illustrating the distribution of predicted potencies across different R2 tail lengths, from LNPs of the top performing head group A5 for the RAW 264.7 cell line. (h) A similar violin plot as in (g), but focusing on LNPs of the entire candidate set. (i) A similar violin plot as in (h), but focusing on R3 tail lengths. (j) A similar violin plot as in (h), but focusing on LNPs of the entire candidate set for Hela cell line. (k, l) Top 2 most important molecular descriptors identified by this model fine-tuned for Hela and RAW 264.7 cell lines respectively, for each head group.

Our subsequent analysis, as illustrated in Fig. 5e, explicates the relationships among lipid candidates targeting Hela cells, as identified by similarities in the AGILE model’s lipid representations. We constructed a similarity network for the chosen 15 lipids, linking each lipid to its nearest equivalents. H9, the most potent LNP, demonstrated connections not only to LNPs with an identical head group (H7, H8) but also to other high-performing candidates, as identified by relative luciferase units (H12, H13). To gain further insights, we carried out molecular explanations on H9, illuminating the most salient regions in the molecule structure that heavily influenced the graph encoder’s prediction within the AGILE model (Fig. 5c, Methods 1.8). Interestingly, head group structures emerged as the most salient for H9, which aligns with our previous findings emphasizing the importance of head groups. Similarly, we developed a similarity network for the 15 candidates selected for RAW 264.7 (Fig. 5f). R6 exhibited connections with other high-performing candidates, including R3, R8, and R11. These four lipids share identical tail structures: one being a C-12 alkyl chain and the other a C-18 alkyl chain. This shared characteristic suggests a strong correlation between these tail structures and the high transfection potency of R6, R3, R8, and R11. Interestingly, both tails are non-biodegradable, which hints at the potential necessity of lipid stability for successful macrophage transfection. Furthermore, these high-performing lipids commonly feature asymmetrical alkyl chains, a trait shared with SM102, which facilitates the formation of an inverted cone geometry more readily ^51^. Similar to the findings for H9, head group structures were identified as an influential factor on the saliency map for R6. Additionally, the tail end was also highlighted as a salient region (Fig. 5d).

Moreover, our results highlight the importance of the carbon chain length of R2 as a critical factor in predicting transfection potency, particularly concerning RAW 264.7. It presents the distribution of predicted potencies relative to varying carbon chain lengths of R2 for lipids hailing from the top-performing head group A5 (Fig. 5g). Two distinct aspects emerge from this distribution: (1) As the R2 carbon chain length increases from 10 to 12, a corresponding uptick in predicted potency becomes apparent. Interestingly, any further extension in the R2 length inversely impacts the predicted potency. (2) In addition, R2’s shorter carbon chain lengths (C≤12) correlate with less variance in potency predictions compared to their longer counterparts (C >12). This trend is not restricted to the top-performing head groups but resonates across others, as well (Supplementary Fig. S14), a phenomenon further corroborated by a Pearson Correlation coefficient of −0.58. Examining the distribution plot for all lipids in the candidate set (Fig. 5h) reveals a similar pattern concerning R2 carbon chain length and predicted potency, albeit with a slightly attenuated Pearson Correlation of −0.39. Notably, we observe less variability amongst the shorter R2 chains (C≤12). Interestingly, the importance of carbon chain lengths varies asymmetrically between the two respective tails. As shown in Fig. 5i, the correlation between the predicted potencies and R3 carbon chain lengths is noticeably lower than that of the R2 carbon chain lengths (−0.15 vs. −0.39). These tail-length findings pertain specifically to transfection in RAW 264.7. As displayed in Fig. 5j, the pattern within the Hela cell line is less defined, resulting in a Pearson correlation of −0.22. Collectively, these insights hold significant implications for guiding the design of LNPs specifically tailored for RAW 264.7.

## Discussion

In this work, we presented the AGILE platform trained on comprehensive virtual and wet-lab libraries to enable predictions of LNP potency across different cell lines even in data-limited settings. Through exposure to an extensive array of molecular descriptors during the training process, the deep learning component in AGILE gained fundamental insights into the complex dynamics of LNP design, incorporating features like electronic and steric properties, and carbon chain lengths in a completely self-supervised manner.

One of the important findings is the influence of the molecular descriptor VSA_EState3 and SssNH on the potency prediction for the Hela cell line. These descriptors, which quantify the electronic and steric properties of a molecule’s surface area and represent a tertiary amine, respectively, align with current expert understanding in lipid design. The connection between these molecular characteristics and their influence on delivery potency exemplifies the power of deep learning in elucidating nuanced molecular features. This correlation between expert knowledge and model interpretation endorses the validity of AGILE’s predictive capabilities and lays a groundwork for future studies on other cell lines. Contrastingly, for the RAW 264.7 cell line, SpDiam_Dzi and VR3_D were identified as the most influential descriptors, highlighting the different physicochemical properties favored by different cell types. This variance in influential descriptors underscores the need for cell-specific LNP design, emphasizing the limitations in applying a one-size-fits-all approach to LNP design across diverse cell lines.

The molecular explanation applied to H9, the most potent LNP for the Hela cell line, further corroborated the importance of head groups, a knowledge already prevalent in LNP design. On the other hand, for RAW 264.7, the high-performing LNPs shared identical tail structures, hinting at the potential role of tail structures in macrophage transfection. The fact that these tail structures are non-biodegradable also implies the significance of lipid stability in LNP potency. Such findings, which would be otherwise elusive without AGILE, elucidate the inherent complexities involved in tailoring LNPs for individual cell types.

Moreover, we found that the carbon chain length of R2 was a critical determinant of transfection potency, particularly in RAW 264.7. This result brings attention to the need for a delicate balance in the chain lengths to achieve optimal transfection, further complicating the LNP design process. The variance in the correlation between predicted potencies and carbon chain lengths for different tails - R2 and R3, as well as the asymmetric importance between the two respective tails, reinforces the idea that LNP design is a delicate process involving numerous factors and dependencies.

Furthermore, we found that AGILE’s predictive power consistently improved with training on larger and more diverse datasets, mirroring observations in fields like natural language understanding, computer vision, and mathematical problem-solving. The exposure to extensive datasets during training also seemed to enhance AGILE’s robustness to various factors and dependencies involved in LNP design. These findings suggest that as we continue to expand our dataset, future models pretrained on even larger scales may yield more precise predictions in elusive tasks with increasingly limited task-specific data. For example, beyond using AGILE to discover LNPs for mRNA delivery to previously unexplored tissues and cell types, there’s an opportunity to expand the wet-lab mRNA transfection data from cell cultures to in vivo data from animal studies and ex vivo data in human tissues. This could potentially boost the efficiency and reliability of LNPs discovered by AGILE for in vivo mRNA delivery in human patients, thereby supporting the clinical development of mRNA LNP products. Additionally, incorporating more diverse combinatorial chemistry methods, along with comprehensive wet-lab data, could further enhance the chemical diversity for AGILE model training^52^. This could allow AGILE to identify ionizable lipids with specific functionalities, such as immunostimulatory properties, essential for mRNA vaccine delivery and cancer immunotherapy. Furthermore, AGILE could adopt recent generative models, like diffusion networks^53, 54^,, to generate *de novo* lipid molecules for specific applications.

Overall, AGILE synergizes the strength of combinatorial chemistry and deep learning, elucidating the intricate dynamics of LNP design and making this insight accessible for a multitude of downstream applications. AGILE’s ability to identify and interpret influential molecular descriptors represents a significant leap forward in the field of nanomedicine, particularly in lipid design. Its capacity in predicting the transfection efficacy of LNPs in diverse cell lines, including challenging ones like macrophages, holds promise for not only improving mRNA delivery but also for guiding CAR cell therapy and other immunotherapeutic strategies. It can potentially accelerate the discovery of potent LNPs and facilitate the design of tailored ionizable lipids for mRNA delivery, thereby contributing significantly to the continuous development of mRNA-based therapeutics and their deployment in clinical settings.

## Materials and methods

Extended materials and methods are available in the supplementary information (SI).

### 1.1 Data Preparation

#### Virtual Library

We utilized Ugi combinatorial chemistry method to design diverse head groups, connecting groups, and two distinct alkyl chains. To be specific, we used the Markush Editor in the ChemAxon Marvin Suite (Marvin 23.4.0, ChemAxon, https://www.chemaxon.com). The resulting virtual library contained approximately 60,000 lipid structures which were then exported into SMILES strings. This virtual library compromises multiple carbon chains, from C6 to C26. In addition, the presence or absence of ester bonds and their position in the carbon chain are used to improve the chemical diversity of the virtual library. The surface charge of LNP is usually determined by the lipids’ head groups. In addition, the head group is critical for mRNA binding. Amine groups are commonly used as lipids’ head groups to form hydrogen bonds with mRNA, especially those containing tertiary amine.

#### Experimental library

Our experimental library contains 20 head groups, 12 carbon chains with ester bonds, and 5 carbon chains with isocyanide head groups. We selected 1200 lipids for chemical synthesis and *in vitro* transfection potency experiments in Hela and RAW 264.7 cell lines. We label the corresponding mRNA transfection potency in cells to each compound for the 1,200 lipids library. And these data are generated by ChemAxon Marvin Suite into SMILE files (SMILE files in SI).

#### Candidate library

The final library used for model prediction is a filtered subset of the virtual library. The filtering contains three steps based on availability and rationality. First, we retained the lipids containing tertiary amine structures. Second, we removed tail chains that were too long (>C18) or too short (<C10) based on expert knowledge of plausible ionizable lipid design ^36^. Last, we select only those reagents commercially available for further validation of the model. Upon completion of the filtering process, the final candidate library comprises approximately 12,000 lipids (SMILE files in SI), with 22 unique head groups (Supplementary Fig. S17), and a distinct arrangement of 9 R2 tail types alongside 2 R3 tail types (Supplementary Fig. S18). In the prediction step of the platform, the model proposed the most promising lipids by predicting and ranking on the candidate library.

### 1.2 Molecular graph construction

Molecular structures can be naturally represented as graphs where atoms are nodes and bonds are edges. For each molecule, the SMILES representation is converted into a molecular graph using RDKit ^55^, and later input to the neural network model in the platform. This representation captures the topological structure and properties of a molecule effectively. An LNP molecule graph *G* is defined as *G* = (*V*, *E*), where nodes *V* represent the atoms and edges *E* represent chemical bonds. The atom node features include the atom type (as on the periodic table) and a flag indicating whether the whole molecule it belongs to is chiral. For a node *v*, the features are constructed in a two-dimensional vector, *ℎ_v_* ∈ *N*^2^. Edge features are constructed based on respective chemical bond types (i.e., single, double, triple, or aromatic bonds) and the stereochemical directionality (i.e., the rdchem.BondDir in RDKit. Similarly, the edge features form another two-dimensional vector for each bond between atom *v* and *u*, *ϵ_v,u_* ∈ *N*^2^.

### 1.3 The Model Architecture

The deep learning model in AGILE comprises three major components: (1) The embedding layers to project node and edge features into learnable vectors, (2) the graph encoder for modeling molecular structures, and (3) the descriptor encoder for modeling molecular properties.

#### Embedding Layers

The embeddings layers project the integer features in ℎ*_v_* and *ϵ_v,s_* to learnable feature vectors 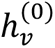 *and* 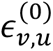, which can be optimized later during the training of the whole neural network. Here, both 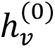 *and* 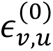 are *R^d^* vectors, and *d* is a predefined size of embedding dimensions. To be specific, we first obtained the embedding vectors for both atom type and charity features in ℎ*_v_*, and added the two vectors elementwise to output the 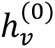:

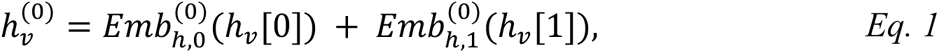

here [i] denotes the i-th element in the vector. *Emb* is the embedding layer projection. In this work, we use the PyTorch Embedding layers (https://pytorch.org/docs/stable/generated/torch.nn.Embedding.html). Similarly, the 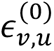 is computed as:

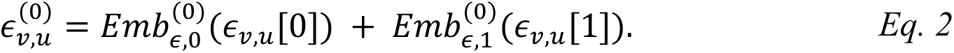

#### Graph Encoder

We used Graph Isomorphism Network (GIN)^56^, a type of graph neural network (GNN), to operate on the input molecule graphs and to learn a representation vector for each LNP molecule. GIN can directly propagate messages among nodes and edges on a graph structure and thus is suitable for processing molecular graphs. Additionally, the advantage of GIN over other GNNs is its ability to distinguish between different graph structures, including isomorphic graphs. This makes GIN more expressive than many other GNNs and a suitable tool for tasks involving molecular graph data. It is worth noting that the implemented GIN model follows the similar structures used in MolCLR^57^, so that we can benefit from the general pretrained molecular model of MolCLR as a warm start for the platform (Methods 1.4). The update rule of GIN for a node representation on the *k^th^* layer is given as:

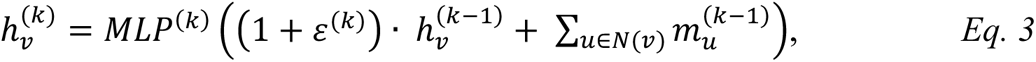

where 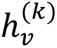 is the representation of node *v* at the *k^th^* layer and *N*(*v*) denotes the set of neighbors of node *v*, and *ε* is a learnable parameter. MLP denotes the stacked fully connected neural network layers. The 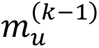is the message propagated between a neighbor *u* to the current node. It is computed as the sum of node and edge contributions:

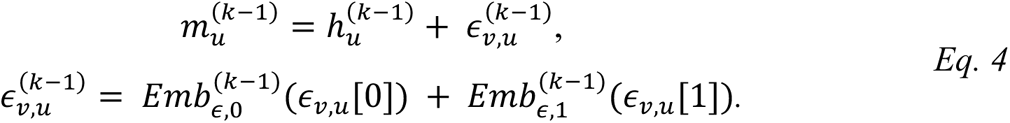

Notably, we use 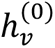 *and* 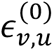 from Eq. 1 and Eq. 2 for the first GIN layer.

We stack a total of *K* GIN layers for the entire Graph Encoder. To extract the feature of the whole molecular graph ℎ*_G_*, we implemented the mean pooling operation on the final layer to integrate all the node features:

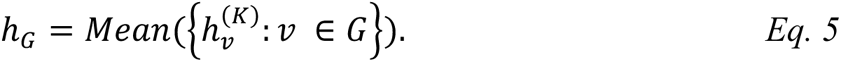

Another fully connected layer is used to transform ℎ_1_ to the final lipid representation *z_G_*:

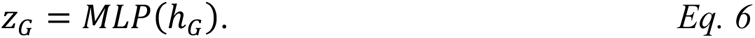

#### Molecular Descriptor Encoder

In addition to the structure features encoded by the GIN, the platform utilizes another descriptor encoder to explicitly model molecular properties. In our experiment, we found this contributes a more stabilized training optimization. We hypothesize that this benefit come from the straight-forward utilization of computed properties during the optimization, which relieves the model from learning all information from the structure alone. In the implementation of the platform, the molecular descriptors derived from Mordred^30^ calculations were used, which contain over 1,000 common descriptors for each molecule, including the num of atoms, num of bonds, et. al. These features are encoded by gully connected layers into a representation for these properties, 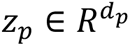:

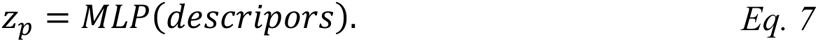

The final representation of the molecule is the concatenation of the structure and property representations:

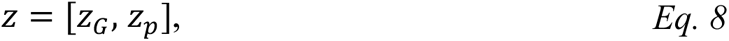

where [,] denotes the concatenation of two vectors.

### 1.4 Model Pre-training

The model pre-training aims to learn generalizable lipid representation that can benefit the downstream transfection potency prediction task. Before our lipid-oriented pre-training, we first initialized the model parameters by the general pre-trained model from MolCLR^57^, which has been trained on over 10 million distinct small molecules. The rationale for this initialization is to provide a warm start of a model that already has been trained to capture molecular structures. Next, we perform continuous pre-training on the 60,000 lipids in the virtual library (Methods 1.1) using contrastive learning to optimize the model’s performance within the LNP domain.

#### Contrastive learning objective

Our pre-training objective is to learn LNP representation through contrasting positive data pairs against negative pairs. The model is trained to minimize the following loss:

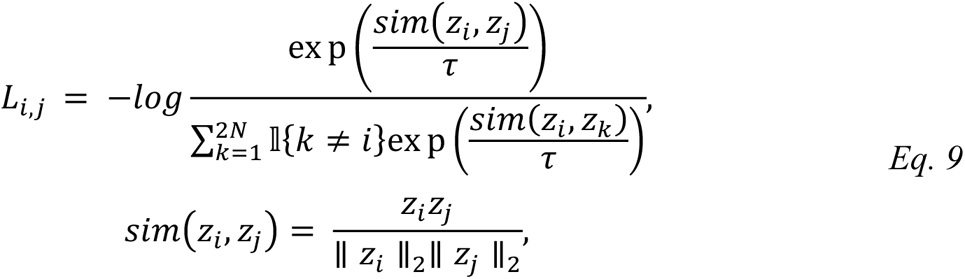

where *z*_*i*_ and *z*_*j*_ are the learned lipid representation vectors extracted from a positive data pair, *N* is the batch size, and *τ* is the temperature parameter set manually. In this pre-training step, we omitted the descriptor encoder, so the lipid representation only contains the graph structure representation *z*_*G*_ as in Eq. 6. To construct the positive data pair, each input lipid molecule graph is transformed into two different but correlated molecule graphs using graph augmentation. The molecule graphs augmented from the same molecule are denoted as a positive pair, and those from different molecules are denoted as negative pairs within each batch. During training, the model learns to maximize the agreement of positive pairs while minimizing the agreement of negative ones.

#### Data Augmentation

We used two augmentation strategies inherited from the MolCLR^57^ pre-training workflow at the atom and bond levels. In the continuous pre-training of LNP molecules, three molecular graph data augmentation strategies are consistently employed. 1) Atom masking: Within the lipid molecular graph, atoms are randomly masked according to a specified ratio. This process compels the model to assimilate chemical information, such as atom types and corresponding chemical bond varieties within lipid molecules. 2) Bond deletion: Chemical bonds interconnecting atoms are randomly removed in accordance with a designated ratio. As the formation and dissociation of chemical bonds dictate the properties of LNP molecules during chemical reactions, bond deletion facilitates the model’s learning of correlations between LNP molecule involvement in various reactions.

### 1.5 Model Fine-tuning

The lipid-oriented pretrained model (Methods 1.4) serves as the starting point of the fine-tuning stage. During the fine-tuning, we included the Molecular Descriptor Encoder and used the combined output *z* in Eq. 8 as the molecule representation. For the property descriptor input, a series of preprocessing procedures are executed, aiming to isolate pertinent features. Initially, descriptors with a standard deviation of zero are eliminated, followed by the selection of descriptors exhibiting correlation with the experimentally determined transfection potency in both Hela and Raw 264.7 cells (R2 score > 0.006), resulting in the identification of 813 salient descriptors (Supplementary Fig. S19). Subsequently, log transformation is applied to descriptors possessing extensive data ranges, with normalization conducted accordingly. The preprocessing steps enacted on the fine-tuning dataset are documented and replicated for the 12,000 lipids in the candidate library in anticipation of the model prediction phase (Methods 1.6).

The model is fine-tuned utilizing the 1,200 lipids of the experiment library to perform regression on LNP transfection potency. The mean squared loss between the predicted and ground-truth potency is used to optimize the model parameters:

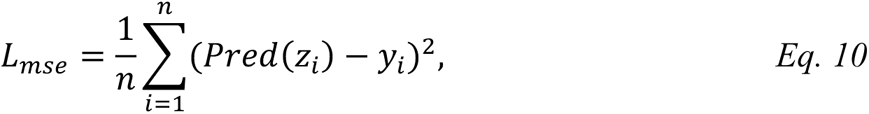

where *Pred*(·) denotes the fully connected layers that perform the potency prediction, and *y*_*i*_ is the actual transfection potency recorded *in vitro*.

A scaffold-based 80%-10%-10% train-valid-test split is performed on the experimental library. We fine-tune the model on the training set only and evaluate the performance on the validation set using root mean squared error (RMSE) and Pearson correlation with the ground truth transfection potency.

### 1.6 Model ensemble prediction and candidate ranking

To enhance the model’s robustness and generalizability, the fine-tuning process is carried out ten times, from which the top five models are selected based on RMSE and Pearson correlation performance on the testing set. These five models are subsequently employed for ensemble prediction on the 12,000-member candidate set. We first get the potency predictions from each model and calculate the average and standard deviation of the five predicted values for each candidate molecule. The mean predicted values are then subtracted from the standard deviation, and the resulting predicted score is used to rank the candidates.

We observed that the predicted potencies exhibit distinct stratification by head groups and tail combinations, and the structural differences among molecules with the same head groups and tail combinations are relatively minor (Supplementary Fig. S20). To increase the diversity of selected candidates, we implement a ranking scheme that sorts candidate LNPs by head groups and tail combinations (Supplementary Fig. S21). Given the predicted values, candidates are first organized by head groups and subsequently ranked in descending order. Candidates within each head group are then ranked by tail combinations following the same schema. Ultimately, we select the top five head groups and the top three tail combinations from each head group, resulting in a final candidate set of 15 LNPs.

### 1.7 Implementation details

The Graph Encoder in the model consists of a five-layer GIN with ReLU activation. To extract a 512-dimensional LNP representation, an average pooling layer is applied to each lipid molecular graph. A single hidden layer MLP is then employed to map the representation into a 256-dimensional latent space. During model pre-training, the contrastive loss is optimized using the Adam optimizer^58^, with a weight decay of 10^.<^, and the temperature is set to 0.1. The pre-training process involves a batch size of 512 for 100 epochs.

For model fine-tuning, an additional MLP with one hidden layer is introduced to map the molecular descriptors into 100-dimensional latent vectors. These vectors are concatenated with the 256-dimensional LNP representation obtained from the GNN encoder. Subsequently, a two-layer MLP is utilized to derive the final prediction value from the concatenated vector. The fine-tuning process employs the Adam optimizer with a weight decay of 10^.=^ to optimize the loss (Eq. 10). Each fine-tuned model is trained using a batch size of 128 for 30 epochs.

### 1.8 Model interpretation

#### Salient molecular descriptors calculation

In our study, we employed the Integrated Gradients^59^ methodology featured in the Captum^60^ Python package to interpret the significance of molecular descriptors. The process involves approximating the integral of molecular descriptor gradients in relation to their respective predicted potencies for each LNP within the candidate library. A molecular descriptor’s prominence is proportionate to the absolute value of its integrated gradient. We implemented computations across all five ensemble models for each target cell line. To calculate an overall significance for each feature, we initially averaged the computed gradients across all input samples on each model, subsequently normalizing these importance scores. The final step involved computing the mean of these importance scores across all five models. The top 20 critical features were selected and visualized based on the calculated importance scores. When assessing feature significance in the context of head groups, we averaged the integrated gradients for each head group and then proceeded to normalization. Following this, we averaged the results across the five models for each respective head group. The top two significant features for each head group were then selected, and their scores were visualized across all head groups.

#### Construction of the similarity network on the selected candidates

We constructed a similarity network for the 15 selected candidates respective to each target cell line, with the aim of elucidating the similarities among the candidates. Utilizing the LNP vector representations provided by the corresponding fine-tuned model, we computed the cosine similarities for each candidate pair and chose the four most similar neighbors for each. This generated similarity network was then visualized, with the node sizes representing the relative luciferase units.

#### Molecular structure interpretation

To ascertain the critical areas within the LNP structure that contribute significantly to the model’s predictions, we engaged the Model Agnostic Counterfactual Compounds Generation feature present in the ExMol Python package^61^. This is accomplished by generating molecular counterfactuals and investigating the alterations required in the LNP molecule to modify its predicted transfection potency (Supplementary Fig. S22). The molecular counterfactuals produced are designed to retain as much similarity to the input LNP molecule as feasible. If modifications in particular regions result in either an increase or decrease in the predicted potency, such areas are deemed as essential regions. The critical areas identified through this process were visualized for both H9 and R6.

### 1.9 Materials and lipid library synthesis

To prepare our materials, we got amines and starting compounds from Sigma-Aldrich and TCI America. We then put 10 µL of a 350 µM stock solution containing amines and tails into each well of a 96-well plate with glass inserts. This stock solution was made by mixing the compounds in a 2:1 ratio of methanol with 0.2 eqv. catalyst phenyl hypophosphoric acid (H_3_PO_4_). The plates were covered and placed on a shaker to stir overnight, with conversions yield typically over 70%. We also formulated lipids into LNP in the same reaction plates. These lipids were purified through flash column chromatography, and their final structures were confirmed using ^1^H 400 MHz NMR spectrometry with CDCl_3_ and tetramethylsilane (TMS) as a standard at UHN Nuclear Magnetic Resonance Core Facility. To further analyze our materials, we obtained high-resolution mass spectra using an LC-Mass spectrophotometer at the Centre for Pharmaceutical Oncology of the University of Toronto.

### 1.10 LNP synthesis and formulation for high throughput screening

To conduct high-throughput screening, we created an organic phase by dissolving a mixture of cationic lipid, DOPE (Avanti), cholesterol (Chol, Sigma-Aldrich), and C14-PEG 2000 (Avanti) in ethanol at a predetermined molar ratio. We prepared the aqueous phase using firefly luciferase mRNA (mLuc, Translate), Cre recombinase mRNA (TriLink BioTechnologies) or EGFP-mRNA (TriLink BioTechnologies) in 10 mM sodium citrate buffer (pH 4.0, Fisher). All mRNAs were stored at −80 °C and were allowed to thaw on ice before use. During the high-throughput screening phase, LNPs were synthesized by mixing an aqueous phase containing the mRNA with an ethanol phase containing the lipids by the OT-2 pipetting robot. The aqueous phase was prepared in a 10 mM citrate buffer with the corresponding mRNA. The ethanol phase was prepared by solubilizing a mixture of ionizable lipid, helper phospholipid (DOTAP, DOPE, cholesterol, and C14-PEG 2000 at pre-determined molar ratios with an ionizable lipid/mRNA weight ratio of 10 to 1.

### 1.11 LNP synthesis and formulation for in vitro and vivo tests

For other *in vitro* and *in vivo* tests, all materials were prepared and processed without nucleases throughout the synthesis and formulation steps.DLin-MC3-DMA and ALC0315 were purchased from Echelon Biosciences. MC3-LNP was prepared at the molar ratio of 50:10:38.5:1.5 (DLin-MC3-DMA:DSPC: cholesterol: DMG-PEG2000) and ALC0315-LNP was prepared at the molar ratio of 46.3:9.4:42.7:1.6 (ALC0315:DSPC: cholesterol: ALC0159 [Echelon Biosciences]). The optimal formulations of H278 and R080 LNPs for the subsequent experiments were determined by the LNP formulation optimization method. Except for the high-throughput screening, the aqueous and ethanol phases were rapidly mixed by pipette at a 3:1 volumetric ratio. Post incubation for 15 min in a 4 °C fridge.

### 1.12 LNP formulation optimization

The statistical software JMP 16 (SAS Institute) analyzed the experimental data. In this Design of experiments (DoE) approach, the four-factor Box-Behnken design was suitable for second-order models comprising 17 preparation runs. The design was cited as a common experimental design for screening crucial factors. In this design, all factors (lipid/mRNA weight ratio, ionizable lipid molar ratio, helper lipid molar ratio, and PEG molar ratio) have low, center, and high levels.

### 1.13 In vitro high throughput screening

The lipid library, which was not purified, was directly combined with ethanol and the aqueous solution of mLuc. For *in vitro* transfection, the lipid-mRNA mixture, containing 0.1 μg of mRNA, was added to pre-seeded Hela and Raw264.7 cells in 96-well plates. Following overnight incubation, the transfection potency of mLuc was measured using the One-Glo Luciferase Assay System (Promega), following the manufacturer’s instructions. The luminescence was quantified using the Cytation imaging reader (BioTek). Finally, the resulting bioluminescence values are assigned to each SMILE string.

### 1.14 In vivo luciferase mRNA for bioluminescence

At 6 h after the intramuscular administration of the mRNA LNPs, mice were injected intraperitoneally with 0.2 ml d-luciferin (10 mg/ml in PBS). The mice were anesthetized in a ventilated anesthesia chamber with 1.5% isofluorane in oxygen and imaged 10 min after the injection with an *in vivo* imaging system (IVIS, PerkinElmer). Luminescence was quantified using the Living Image software (PerkinElmer). C57BL/6 mice (4-8 weeks) were purchased from the Jackson Laboratories.

### 1.15 ROSA*^mT/mG^* Cre reporter mice transfection analysis

All animal studies were approved and conducted in compliance with the University Health Network Animal Resources Centre guidelines. For gene recombinant Cre mRNA delivery, LNPs co-formulated with Cre mRNA (0.5 mg kg ^-1^) were i.m. injected into ROSA*^mT/mG^* Cre reporter mice (The Jackson Laboratory). After 7 d, mice were killed, and major organs were collected and imaged using an IVIS imaging system (PerkinElmer). For direct fluorescence imaging, organs and muscle tissues were fixed in 4% buffered paraformaldehyde overnight at 4°C, then equilibrated in 30% sucrose overnight at 4°C before freezing in OCT. Three nonconsecutive sections from each organ sample were mounted with DAPI to visualize nuclei and imaged for DAPI, tdTomato, and GFP. Sectioned into 10 μm depth, and further imaged using a Fluorescence microscope (Zeiss AXIO Observer 7 Inverted LED Fluorescence Motorized Microscope).

### 1.16 Intracellular delivery of GFP mRNA to RAW 264.7

For GFP mRNA delivery, GFP mRNA LNPs containing 500 ng GFP-mRNA were added to 24-well plates for 48 h incubation at 37 °C. Finally, a fluorescence microscope (Zeiss AXIO Observer 7 Inverted LED Fluorescence Motorized Microscope) was used to evaluate the transfection effect.

### 1.17 Statistical analysis

The data were subjected to statistical analyses using GraphPad Prism 9 (GraphPad Software). A two-tailed unpaired Student’s t-test was conducted to assess the significance of the comparisons as indicated. Data are expressed as mean ± s.d. P values <0.05 (*), P < 0.01 (**), P < 0.001 (***) and P < 0.0001 (****) were statistically significant.

## Supporting information

Supplementary Figures S1-S22, Table S1 and Notes will be used for the link to the file on the preprint site.

## Acknowledgments

The authors are grateful to R.S. Langer for project discussions and constructive input. This work was supported by the Leslie Dan Faculty of Pharmacy startup fund, the Princess Margaret Cancer Center operating fund, the Connaught Fund (no. 514681), the J. P. Bickell Foundation (no. 515159), the Canada Research Chairs Program (no. CRC-2022-00575), Canadian Institutes of Health Research (no. PJH-185722), Natural Sciences and Engineering Research Council of Canada (no. RGPIN-2023-05124) and the Canada Foundation for Innovation - John R. Evans Leaders Fund (no. 43711); Y.X. acknowledge the Postdoctoral Fellowship from PRiME-UHN Clinical Catalyst Program (no. PRMUHN2022-005); A.V. acknowledges the Postdoctoral Fellowship from the PRiME - Precision Medicine initiative at the University of Toronto; R.X.Z.L. acknowledges the Postdoctoral Fellowship from the Acceleration Consortium at the University of Toronto. The authors acknowledge the technical support from the Centre for Pharmaceutical Oncology in Flow Cytometry, and Imaging Facilities, and acknowledge the Princess Margaret Cancer Centre for the use of NMR and Animal facilities. Balloon plots created with bioinformatics.com.cn. Figures 1-4 were created with Biorender.com.

