## Supplementary Figures S1-S22, Table S1 and Notes will be used for the link to the file on the preprint site. for "AGILE Platform: A Deep Learning-Powered Approach to Accelerate LNP Development for mRNA Delivery"

[#] These authors contributed equally.

### Supplementary Figures

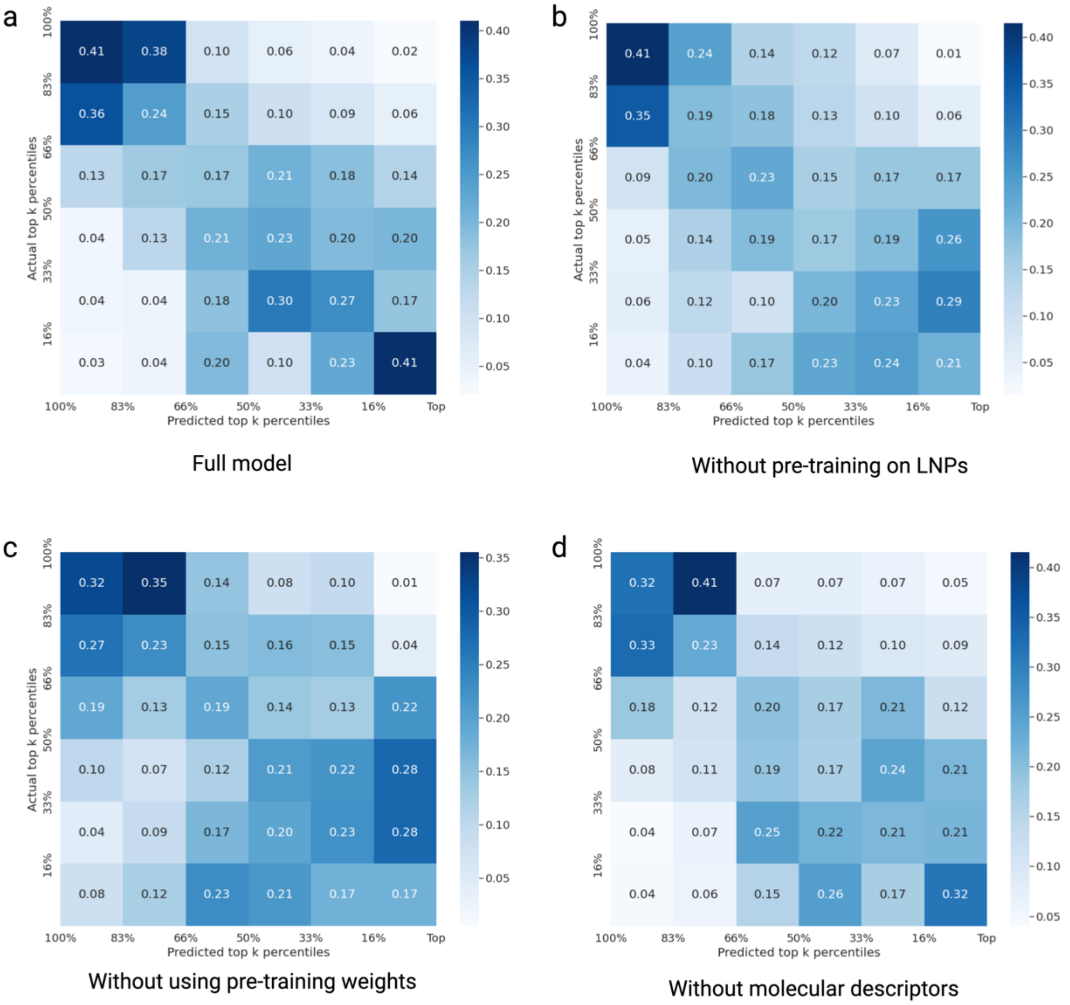


**Figure S1.** This figure provides an ablation study of the AGILE pipeline design, assessing the effect of individual components on overall model performance. (a) Presents the performance of the fully featured model, which has been fine-tuned on the Hela experimental dataset. (b) Depicts the performance of a model variant that does not include pre-training on lipids. Although this iteration demonstrates a comparable ability to identify lower-performing lipids, a notable decrease in its capacity to discern top-performing instances is observed. (c) Showcases the performance of the model when pre-training weights are excluded. In this case, the model's ability to identify top performers decreases further, along with a moderate reduction in its capability to identify lower-performing instances, underlining the significant contribution of pre-training weights to model performance. (d) Highlights the performance of the model when the molecular descriptor module is not incorporated. In this scenario, a concurrent and somewhat significant decline in the model's ability to identify both top and bottom performers is observed suggesting the importance of this module in the overall performance of the AGILE pipeline.

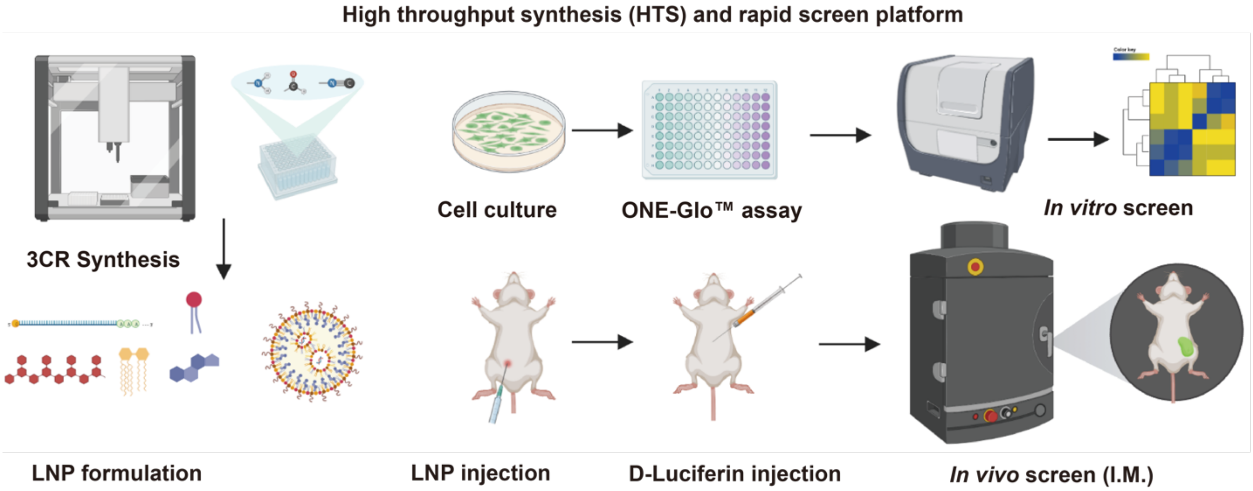


**Figure S2.** Schematic diagram of the high-throughput synthesis and rapid screening platform.

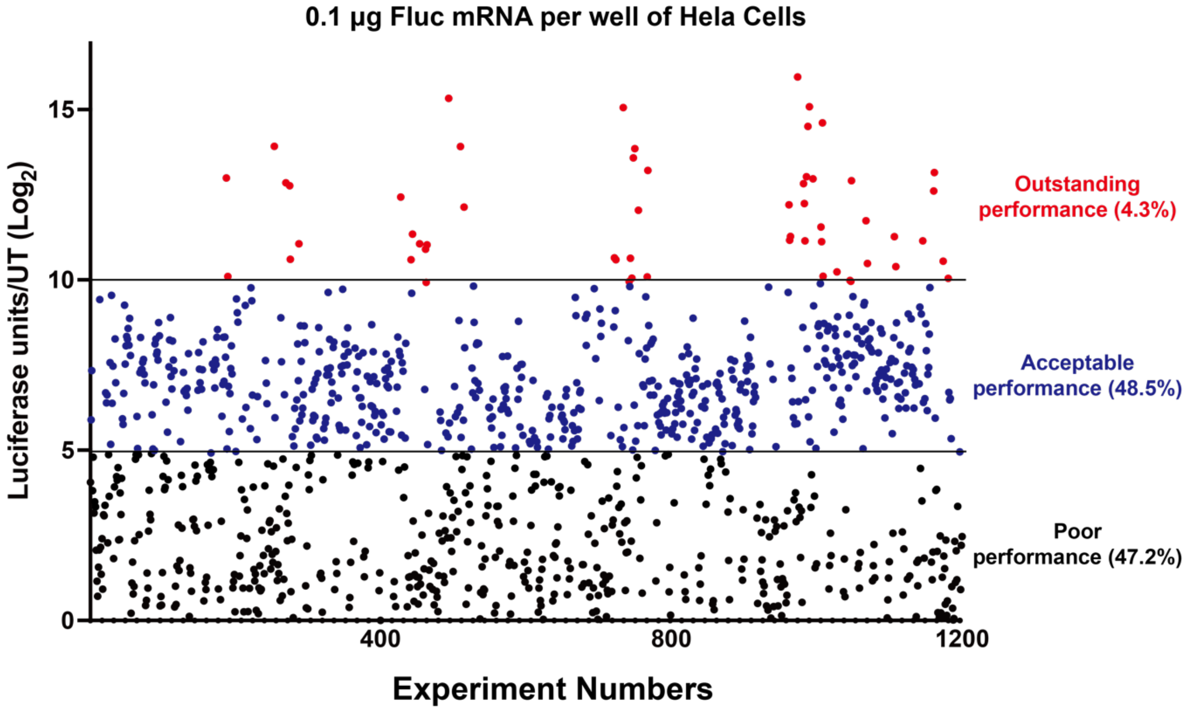


**Figure S3.** Luciferase activity/untreated are shown as scatter plots for the 1,200 LNPs in Hela.

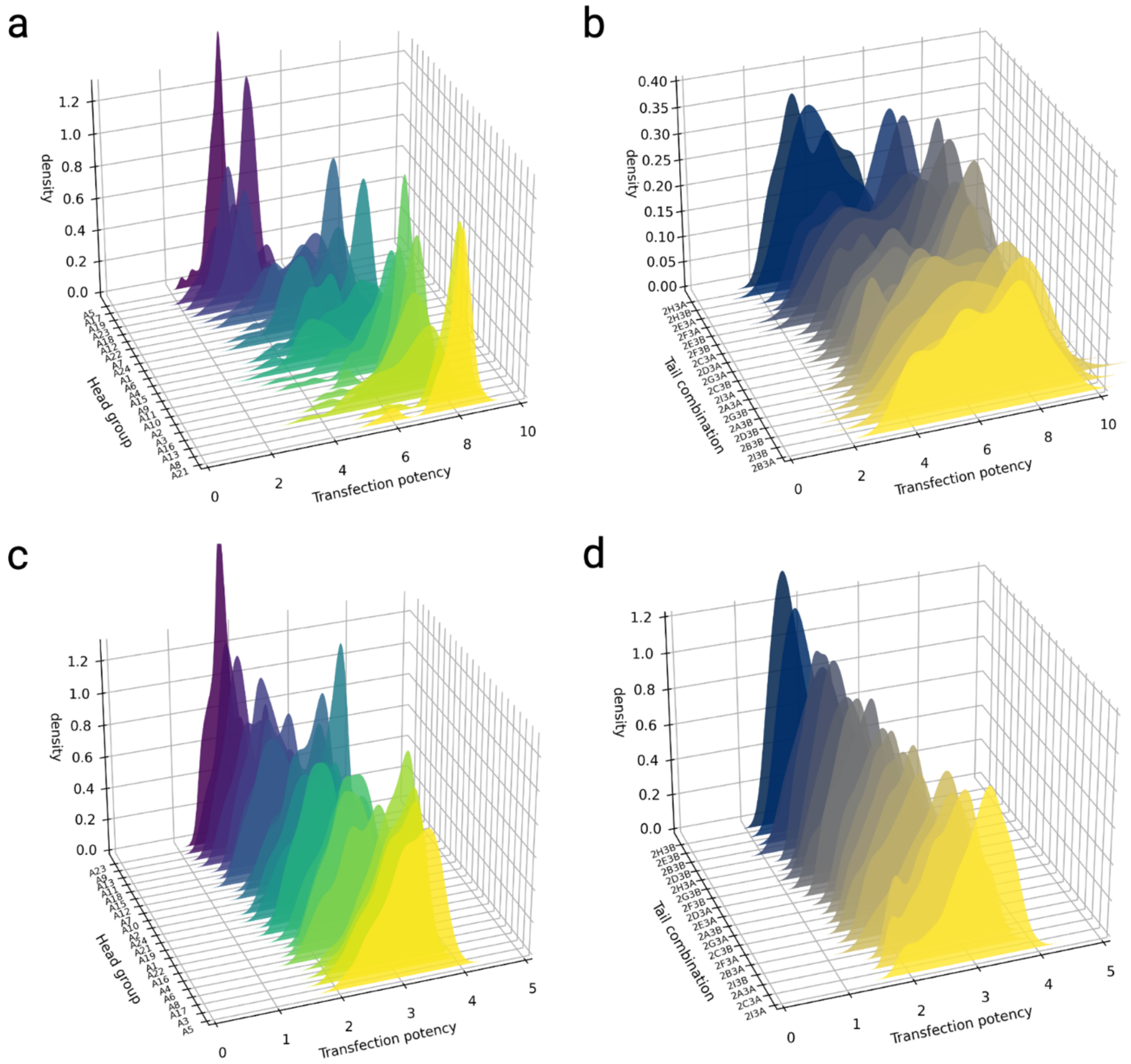


**Figure S4.** The stratified distribution plots for predicted potencies across different categories. (a) Distribution plot demonstrating the potencies predicted for the Hela cell line, stratified according to head group classifications. (b) Similar stratified distribution plot for the Hela cell line, but the stratification is based on tail combinations. (c) Moving to the RAW 264.7 cell line, distribution plot of predicted potencies stratified by head groups. (d) Distribution plot of predicted potencies for the RAW 264.7 cell line, stratified by tail combinations.

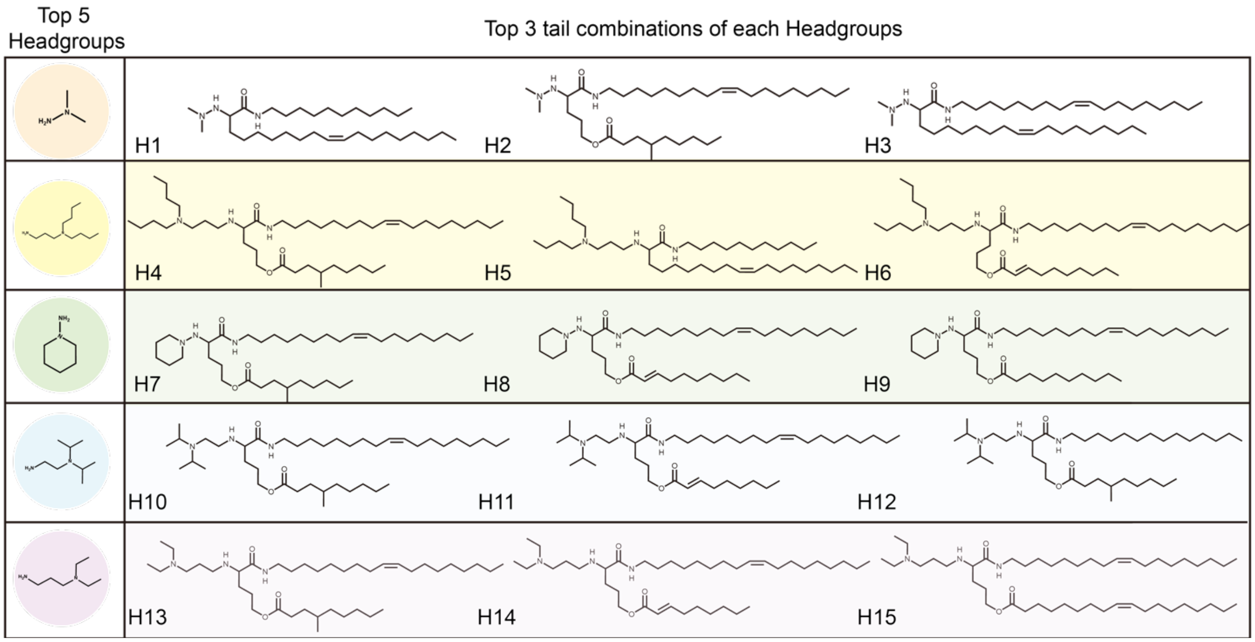


**Figure S5.** Top 15 lipid candidate structures predicted for Hela cells by the AGILE model.

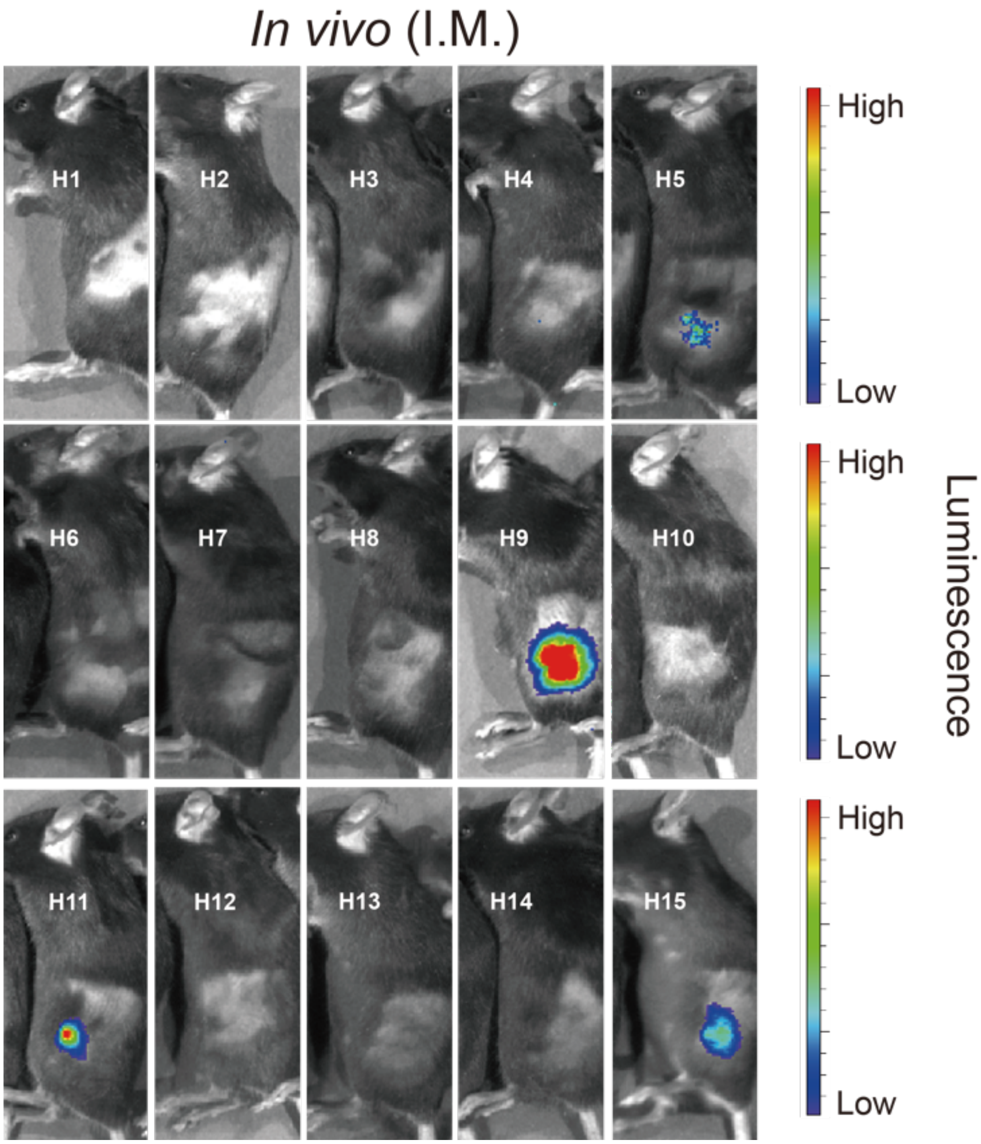


**Figure S6. Ionizable lipids predicted by ML were rapidly synthesized and screened both *in vitro* and *in vivo*.** LNP of unpurified ionizable lipids for intramuscular injection. LNPs formulated with FFL encoding mRNA were injected intramuscularly into mice (0.2 mg mRNA/kg mouse). The FFL expression was visualized at 5 h.

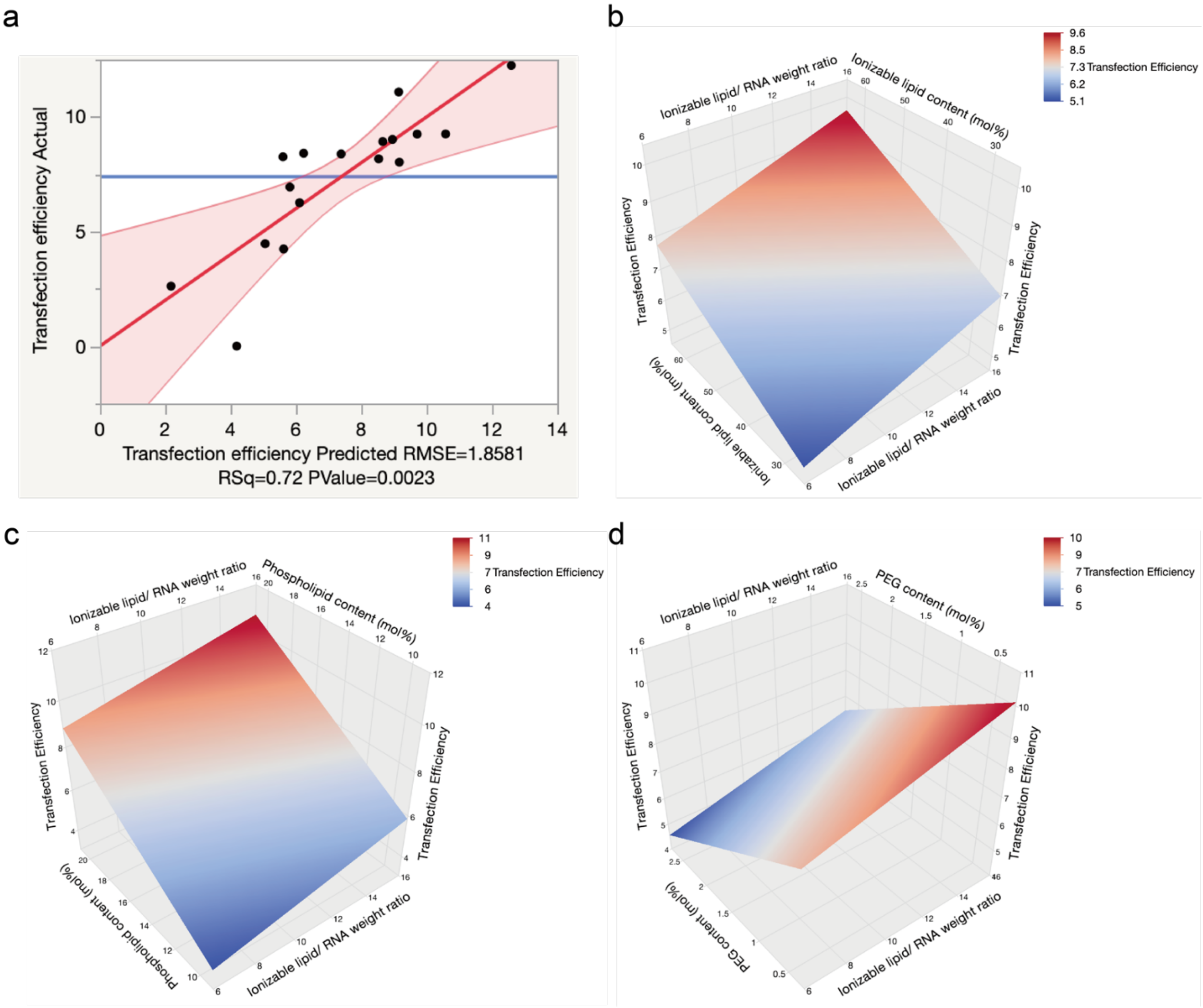


**Figure S7.** LNP formulation optimization and biodistribution by DoE for Hela cells. (a) The calculation for the predicted response for root mean square error (RMSE) in DOE. Different response plots (b) ionizable lipid content (mol%) (c) phospholipid content (mol/%) and (d) PEG content (mol%).

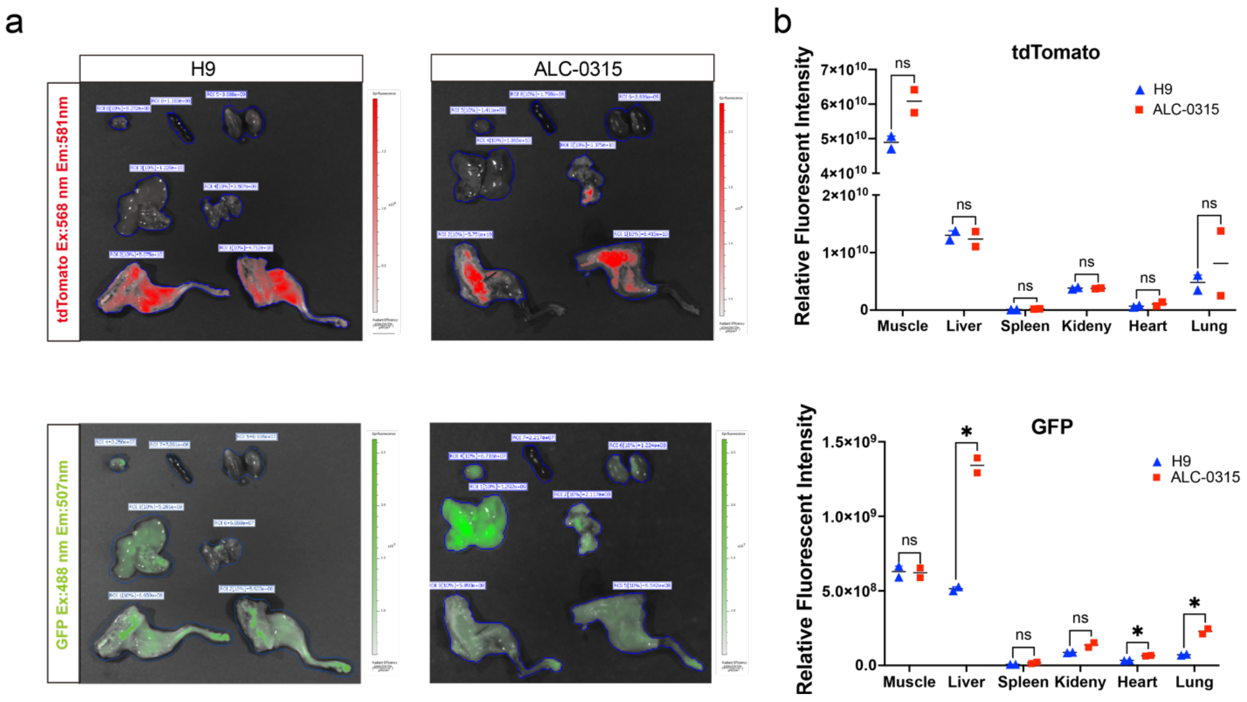


**Figure S8.** (a) IVIS images of ROSA$^{mT/mG}$ Cre reporter mice organs after cre-mRNA LNP by Intramuscular injection. (0.5 mg kg $^{-1}$) (b) Comparison of the relative fluorescence intensity of each organ after intramuscular injection of cre-mRNA LNP. (tdTomato: Ex:568 nm, Em:581 nm, GFP: Ex: 488 nm, Em: 507 nm)

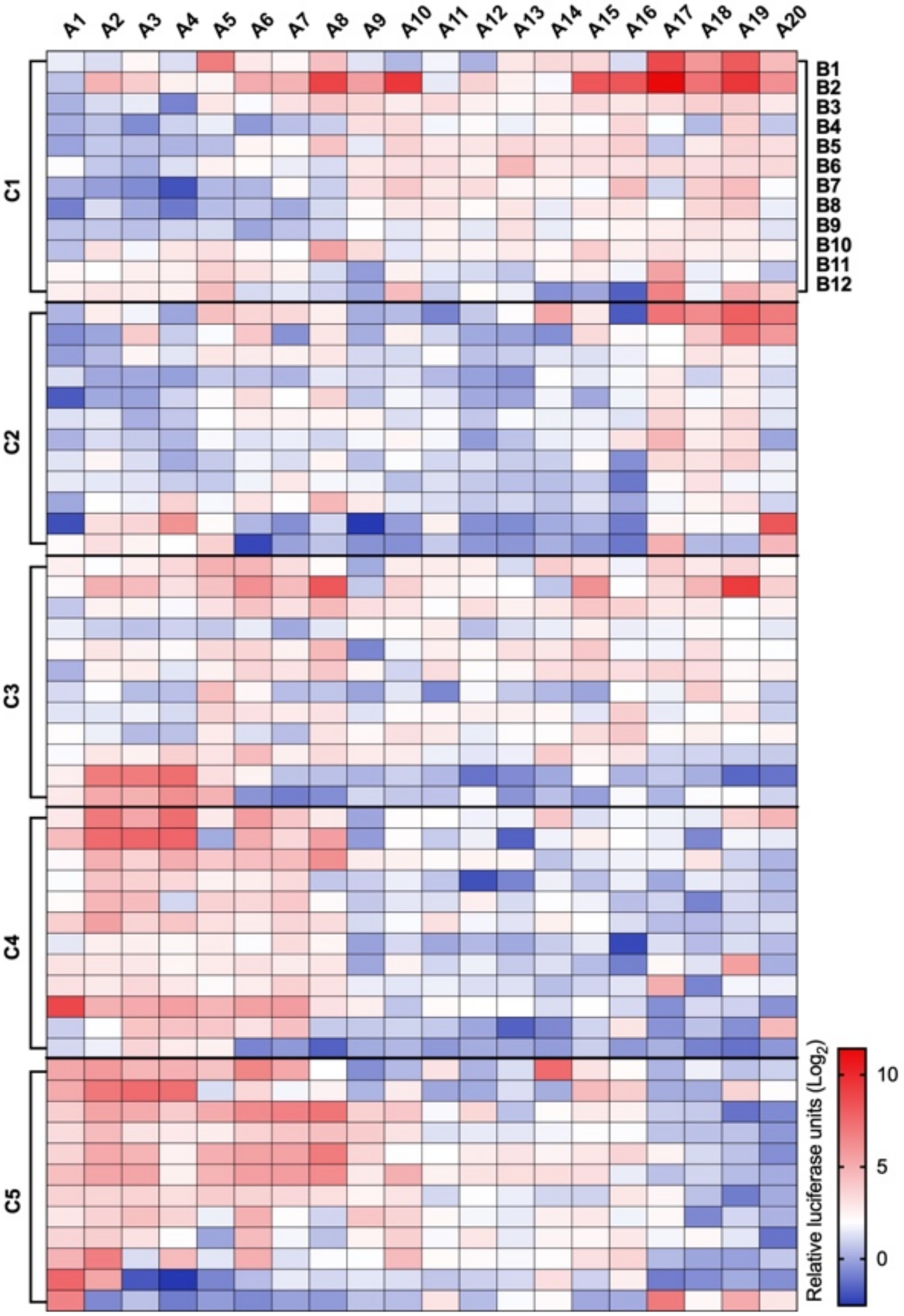


**Figure S9.** The data used for the fine-tuning are depicted in a heatmap, which involved 1,200 LNPs for Fluc mRNA (mLuc) delivery and measuring the relative luciferase expression in RAW 264.7 cells.

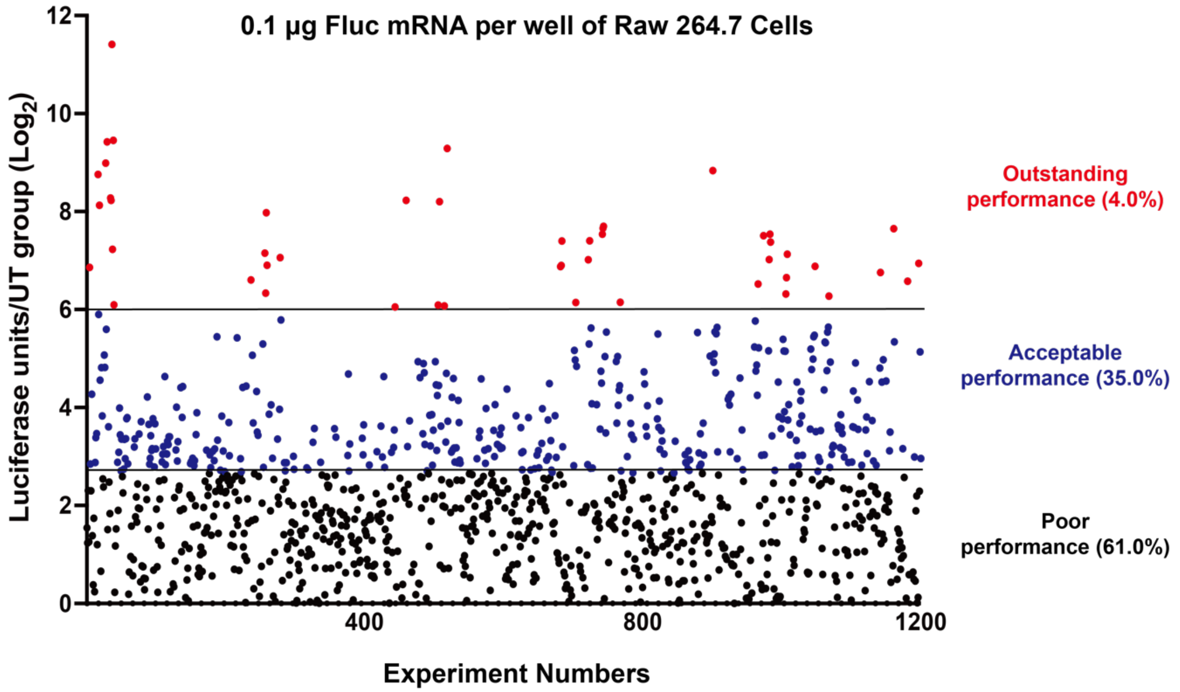


**Figure S10.** Luciferase activity/untreated are shown as scatter plots for the 1,200 LNPs in RAW 264.7.

ML predicted lipid structure for Raw 264.7

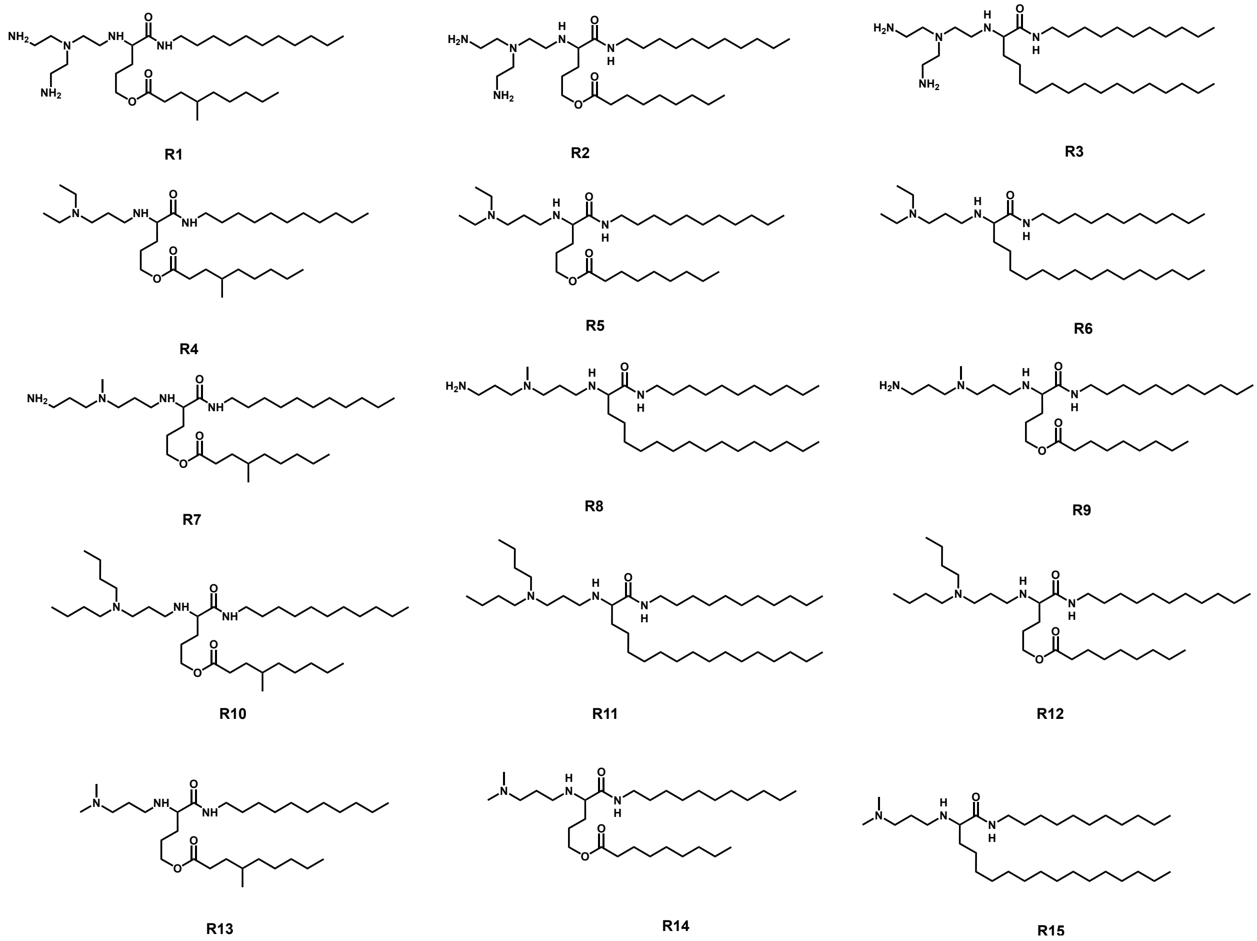


**Figure S11.** Top 15 lipid candidates predicted for RAW 264.7 cells by the model.

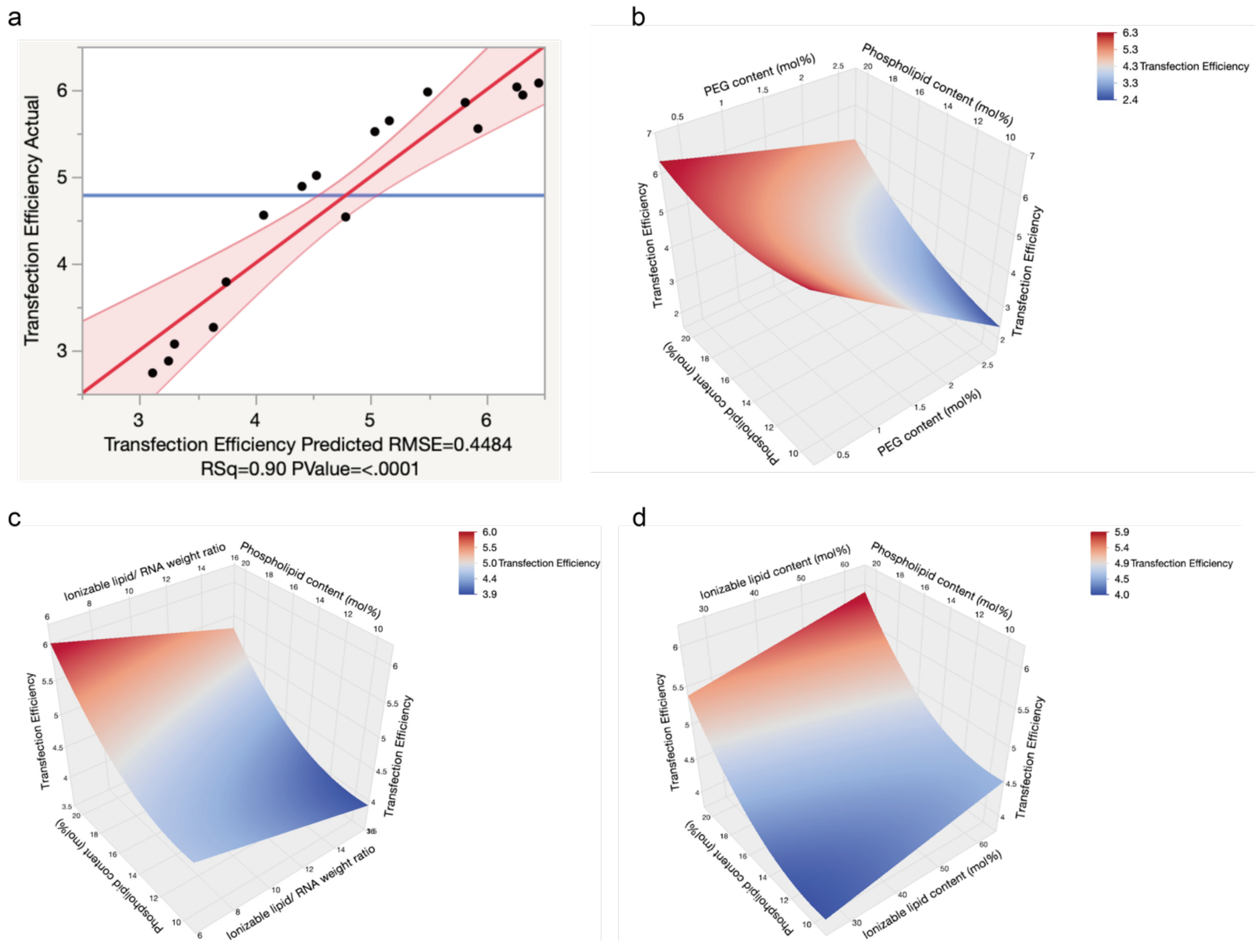


**Figure S12.** LNP formulation optimization and biodistribution by DoE for RAW 264.7 cells. (a) The calculation for the predicted response for root mean square error (RMSE) in DOE. Different response plots (b) PEG content (mol%) (c) phospholipid content (mol/%) and (d) ionizable lipid content (mol%).

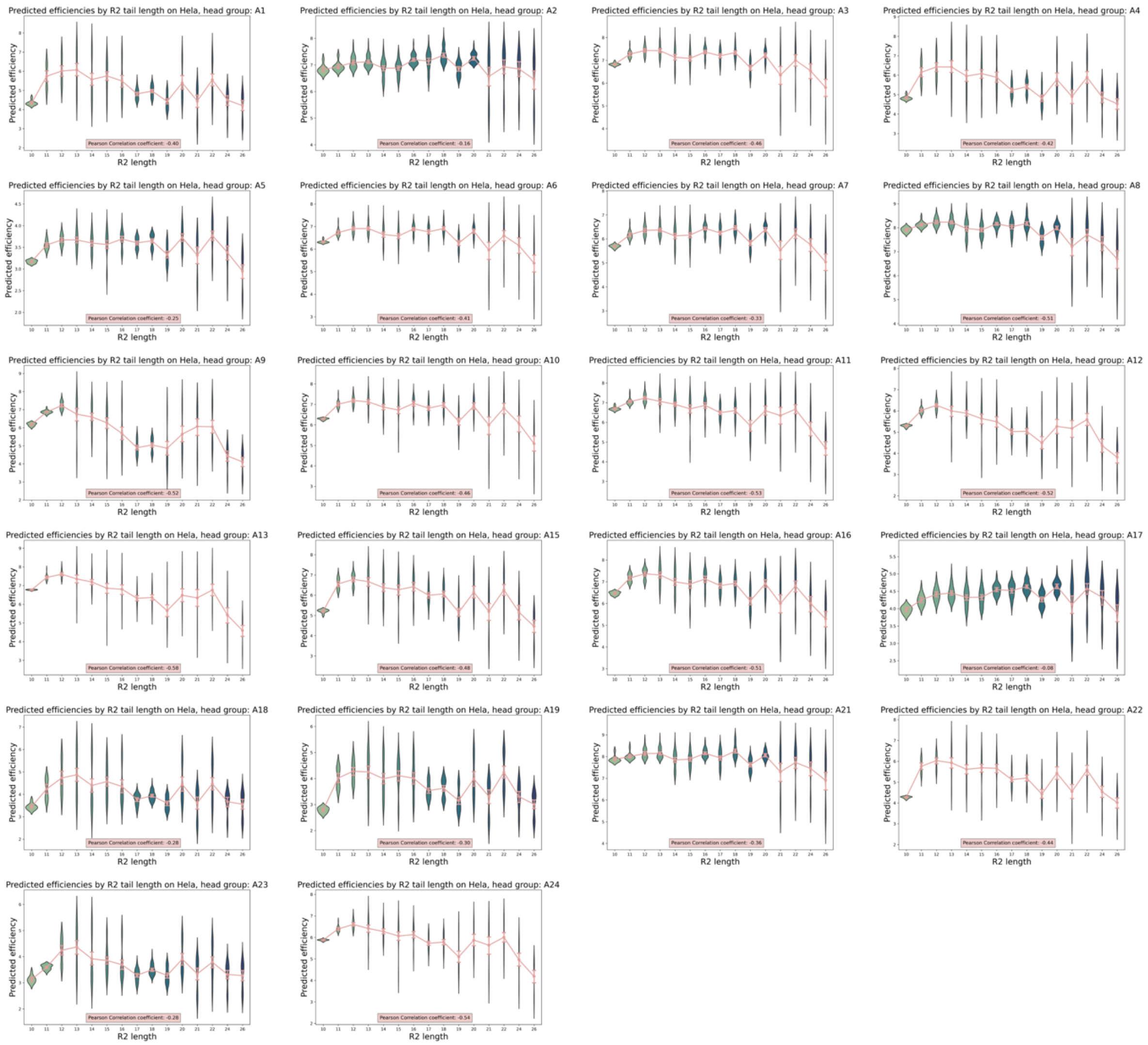


**Figure S13.** Violin plot that visualizes the distribution of predicted potencies for the Hela cell line, categorized based on varying R2 tail lengths from LNPs of each head group. Despite the distinct patterns exhibited by different head groups, a shared trend can be observed: an increase in the R2 carbon chain length from 10 to 12 correlates with an increase in predicted potency. Conversely, any subsequent lengthening of the R2 chain tends to have a detrimental effect on the predicted potencies.

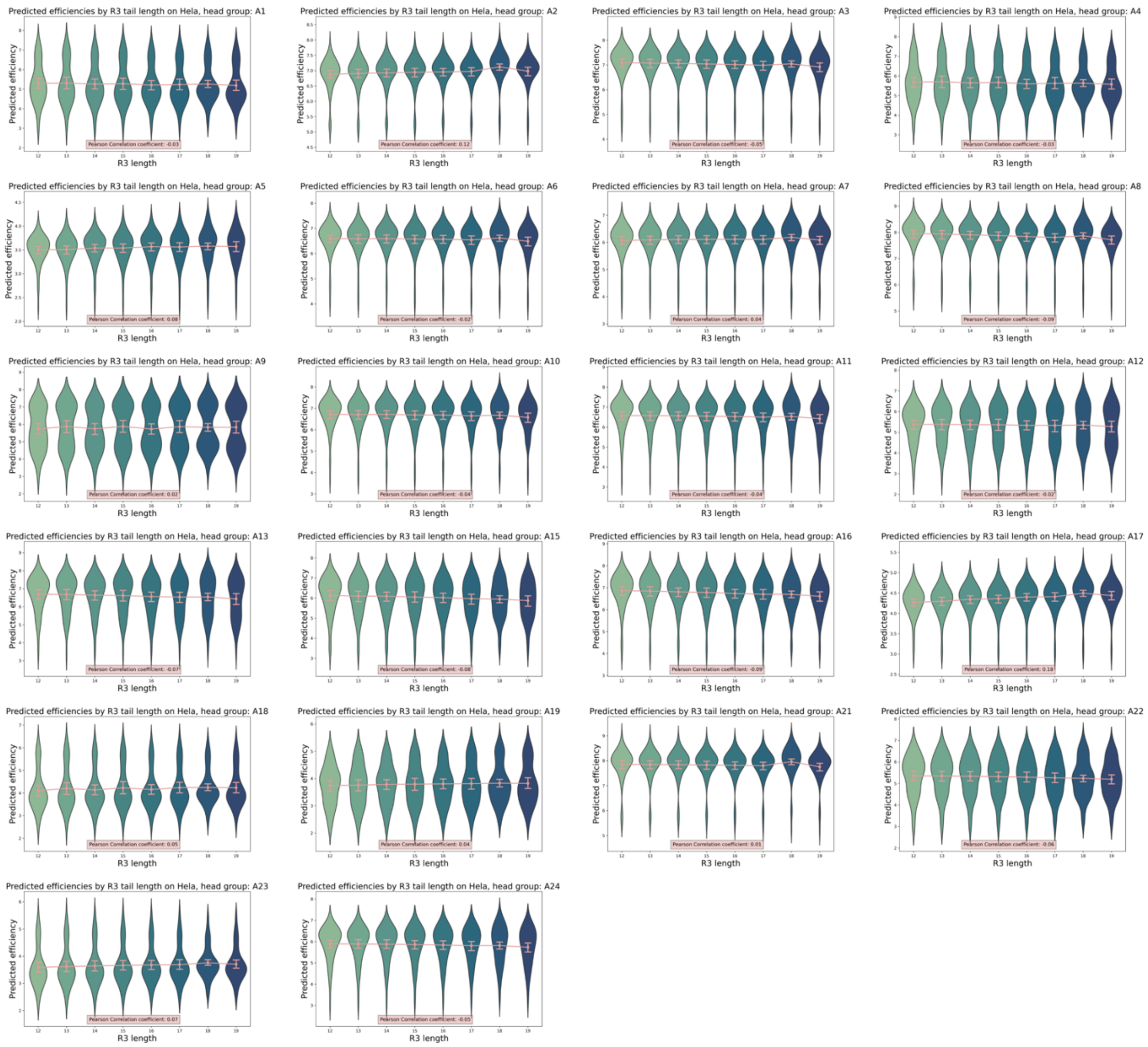


**Figure S14.** Violin plot that visualizes the distribution of predicted potencies for the Hela cell line, categorized based on varying R3 tail lengths from LNPs of each head group. The figure reveals that alterations in R3 tail lengths have a minimal effect on predicted potencies.

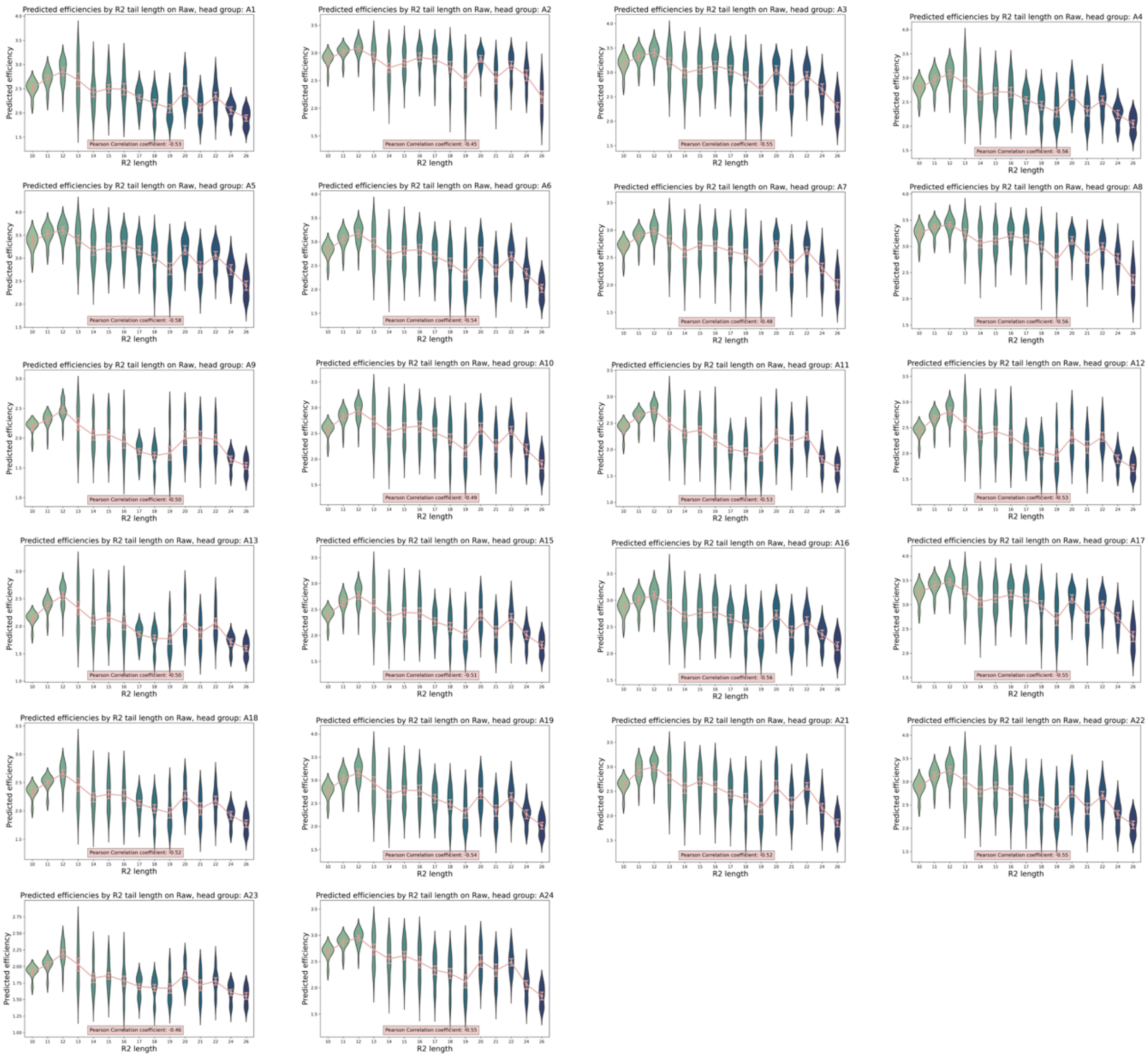


**Figure S15.** Violin plot that visualizes the distribution of predicted potencies for RAW 264.7, categorized based on varying R2 tail lengths from LNPs of each head group. Two shared trends can be observed: (1) An increase in the R2 carbon chain length from 10 to 12 correlates with an increase in predicted potencies. Conversely, any subsequent lengthening of the R2 chain tends to have a detrimental effect on the predicted potencies. (2) shorter carbon chain lengths of R2 (C⩽12) exhibit less fluctuation in potency predictions compared to their longer counterparts (C>12).

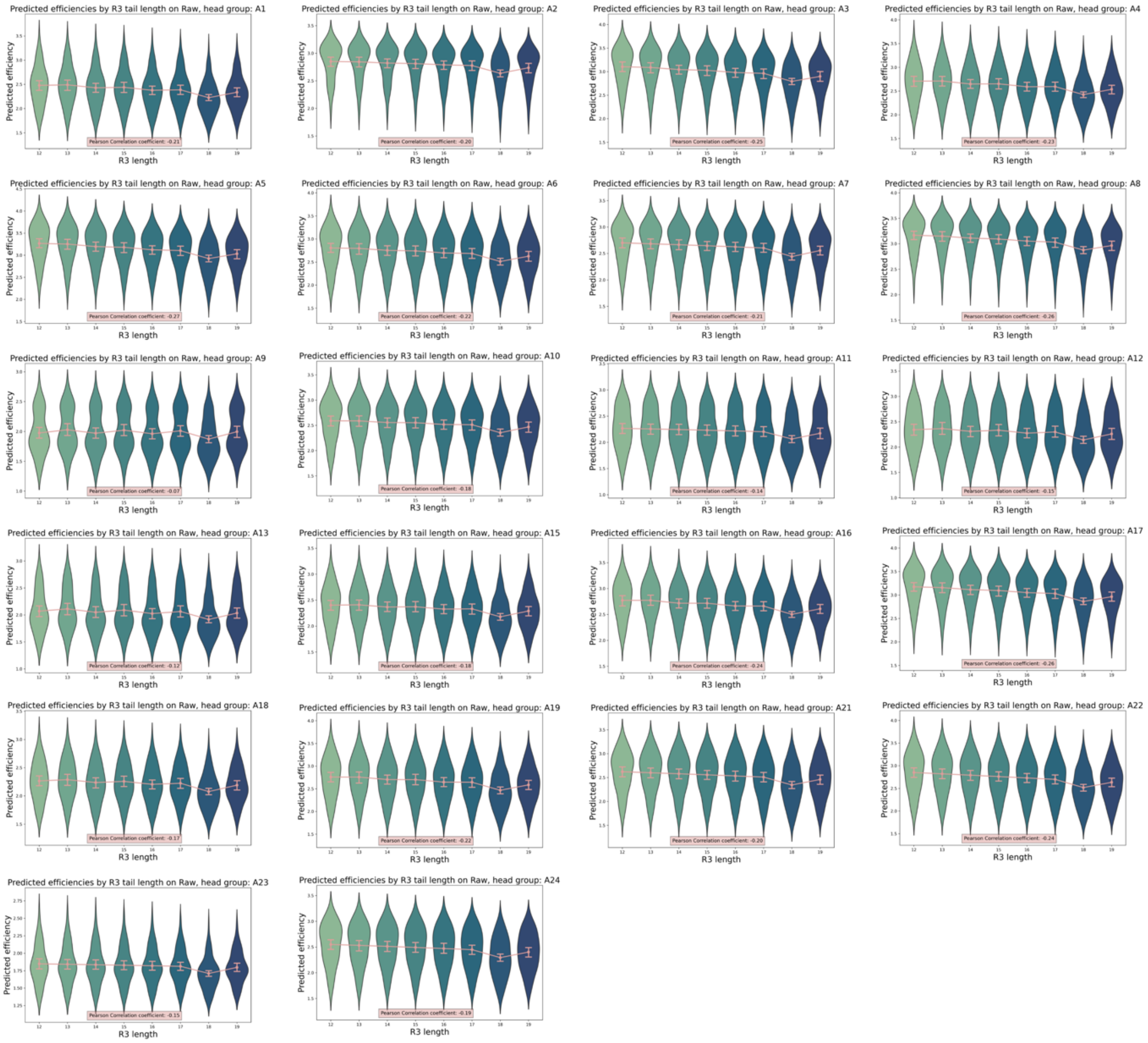


**Figure S16.** Violin plot that visualizes the distribution of predicted potencies for RAW 264.7, categorized based on varying R3 tail lengths from LNPs of each head group. The figures show a shared trend, where smaller R3 tail lengths are associated with higher predicted potencies.

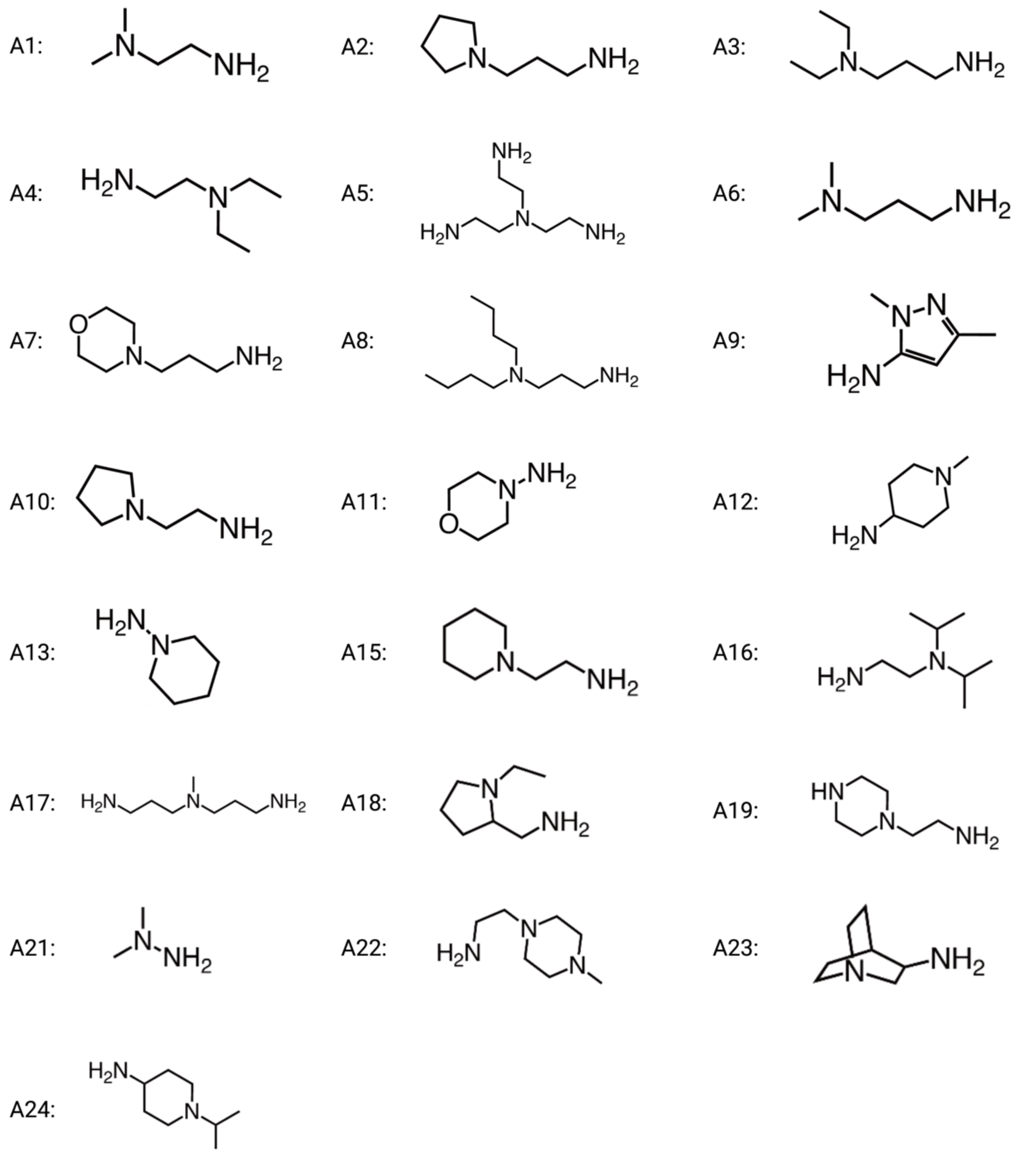


**Figure S17.** Figure displaying the complete collection of 22 distinct head groups incorporated within the candidate library.

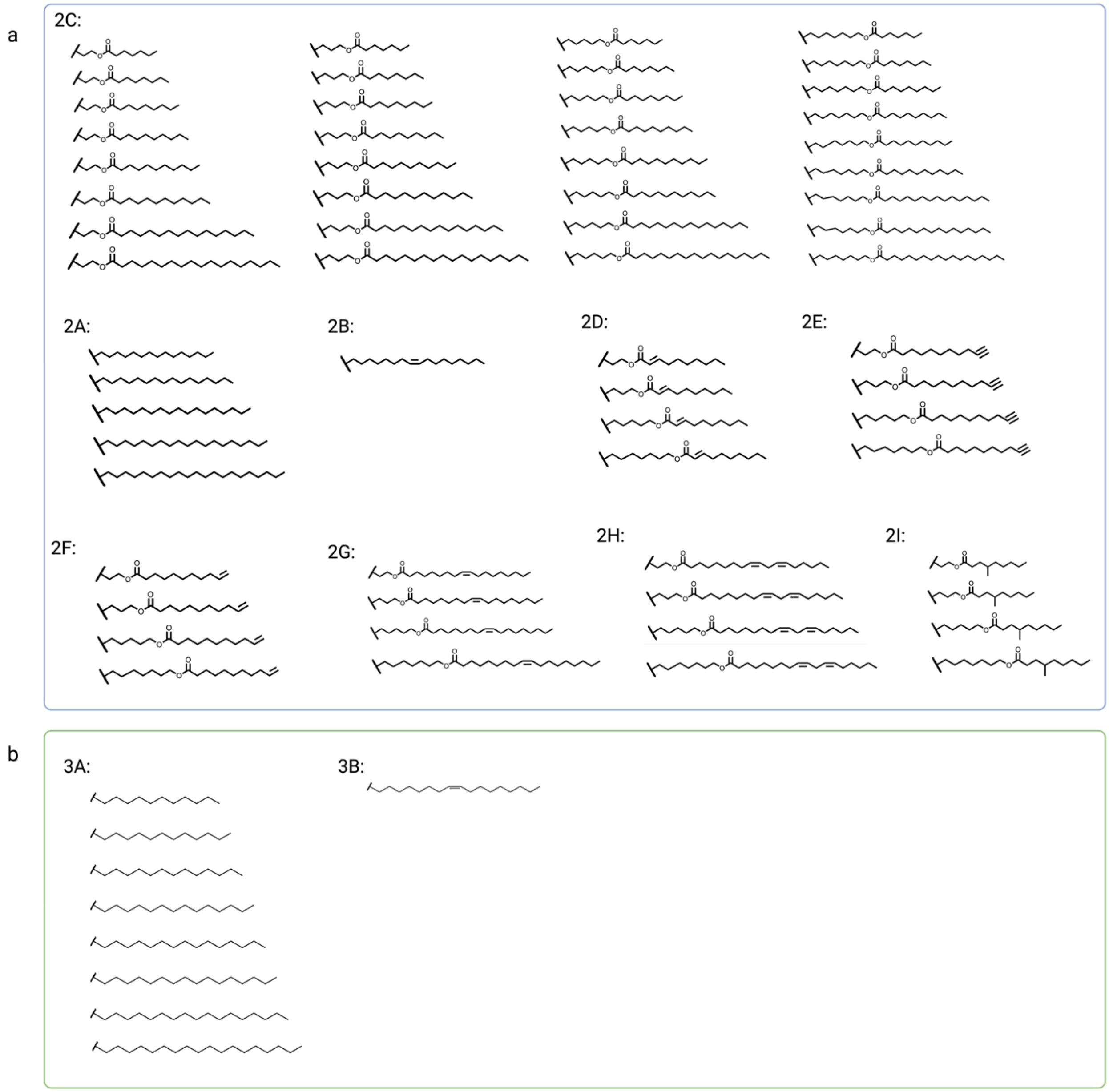


**Figure S18.** Comprehensive representation of all distinct tail combinations present in the candidate library. (a) Showcases different R2 tails. (b) Showcases different R3 tails.

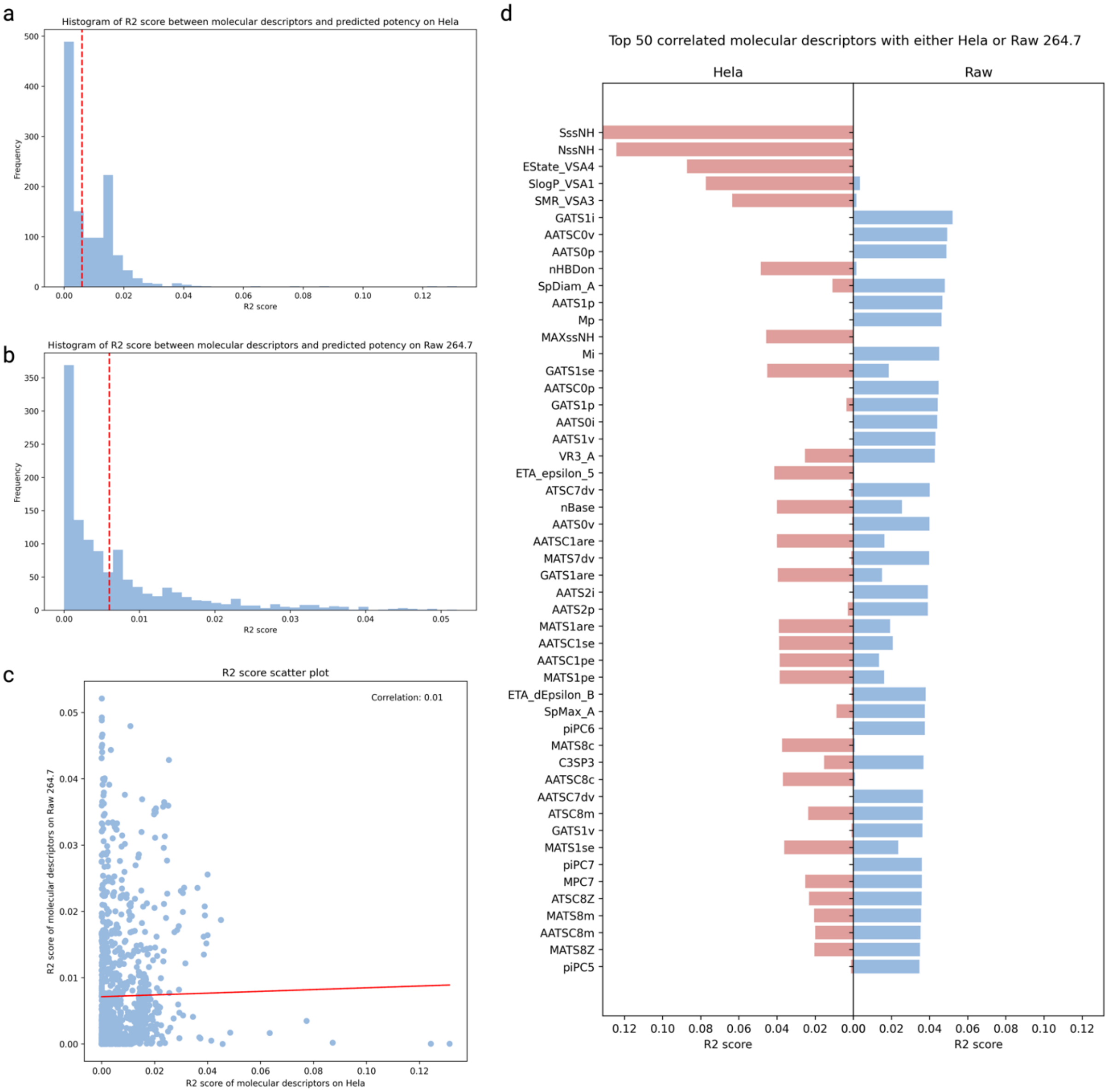


**Figure S19.** (a) A histogram illustrating the R2 score between molecular descriptors and potency labels in the Hela cell line. The red dashed line signifies the 0.006 threshold, established for descriptor selection. (b) Similarly, a histogram is displayed for the R2 score between molecular descriptors and potency labels in the RAW 264.7 cell line, again with the red dashed line denoting the 0.006 threshold for descriptor selection. (c) A scatter plot presents the R2 score of molecular descriptors with potency labels on both Hela and RAW 264.7 cell lines, with values on the two axes. A correlation of 0.01 indicates that molecular descriptors demonstrate a distinct correlation pattern for the Hela cell line and RAW 264.7. (d) Lastly, the top 50 selected molecular descriptors are presented, along with their respective correlation values.

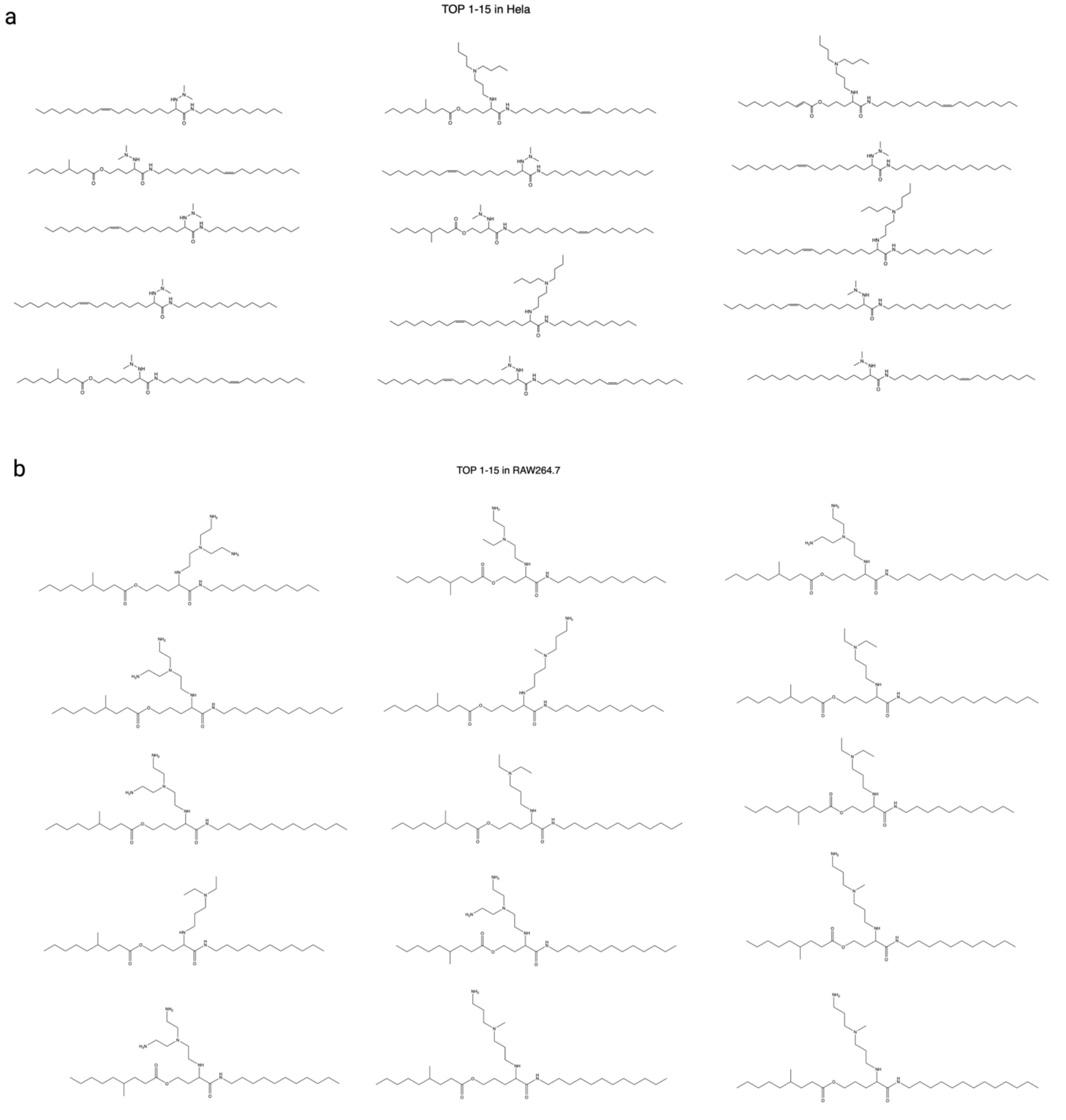


**Figure S20.** Figure illuminating the significance of our head and tail-wise ranking scheme in enhancing structural diversity, exemplified by the top 15 predictions chosen based solely on predicted scores without the application of our ranking method. (a) The top 15 predictions selected for the Hela cell line, without the implementation of our ranking scheme, are displayed. These selections reveal limited head group diversity, as only two different head groups are selected, with 11 being A21 and 4 being A8. (b) The top 15 predictions chosen for the RAW 264.7 cell line, again without our ranking scheme, are shown. These selections also exhibit restricted head group diversity, with only three different head groups represented: 7 are A5, 4 are A3, and 4 are A17.

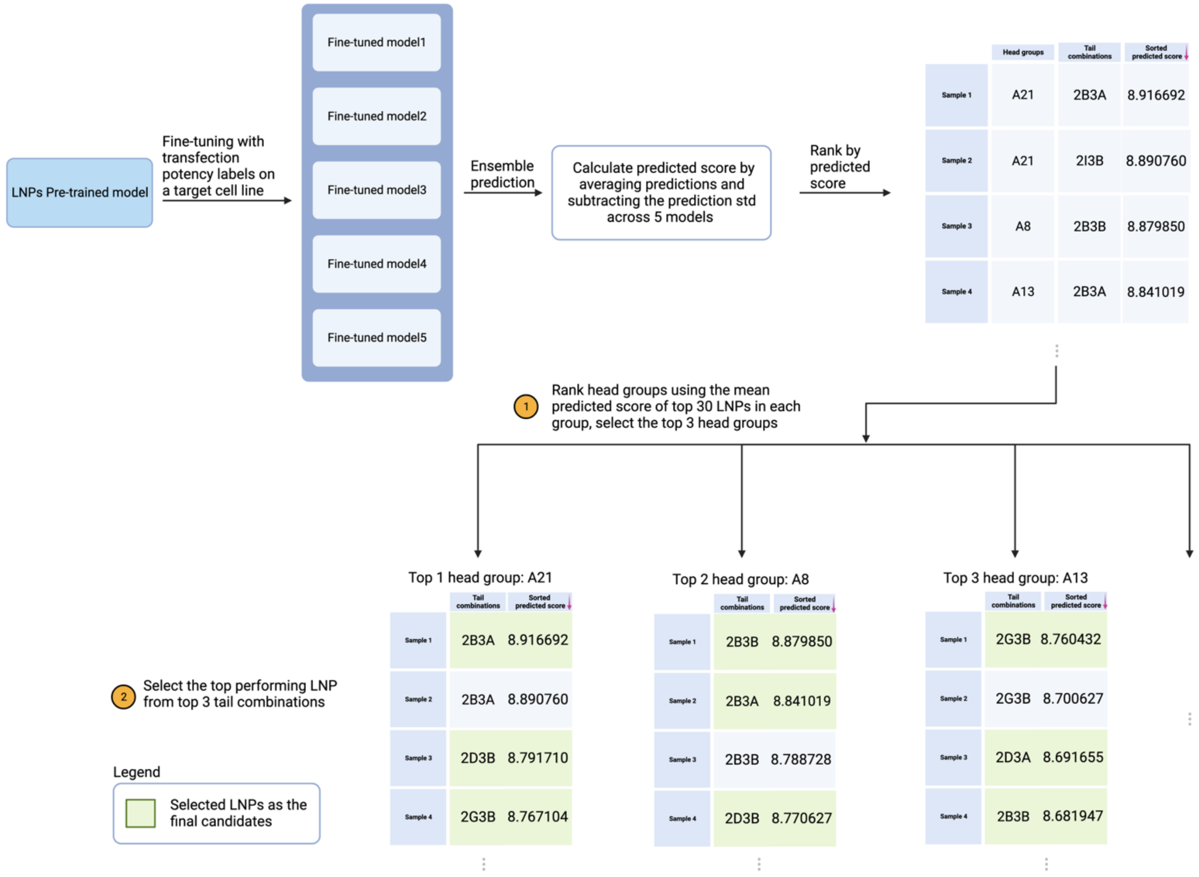


**Figure S21.** This figure provides a comprehensive outline of the ranking and selection procedure implemented by AGILE, exemplified by the selection of the top three head groups and top three tail combinations as the final candidates. Initially, AGILE, pre-trained with lipids, is fine-tuned using measured transfection potency labels for a specific target cell line. The five optimal fine-tuned models are preserved for ensemble prediction. During this ensemble prediction stage, we initially calculate the predicted score by averaging the predictions and then subtracting the standard deviation of the predictions across the five models. This score is subsequently used to rank the samples. Upon obtaining this ranking, a head and tail-wise ranking is conducted to finalize candidate selection. The initial step utilizes the mean predicted score of the top 30 lipids in each group to rank the head groups, enabling the selection of the top three head groups. Subsequently, within these chosen head groups, the highest-performing lipid from the top three tail combinations is selected. This process results in a final selection of nine candidates, comprising three each from the top three head groups and tail combinations.

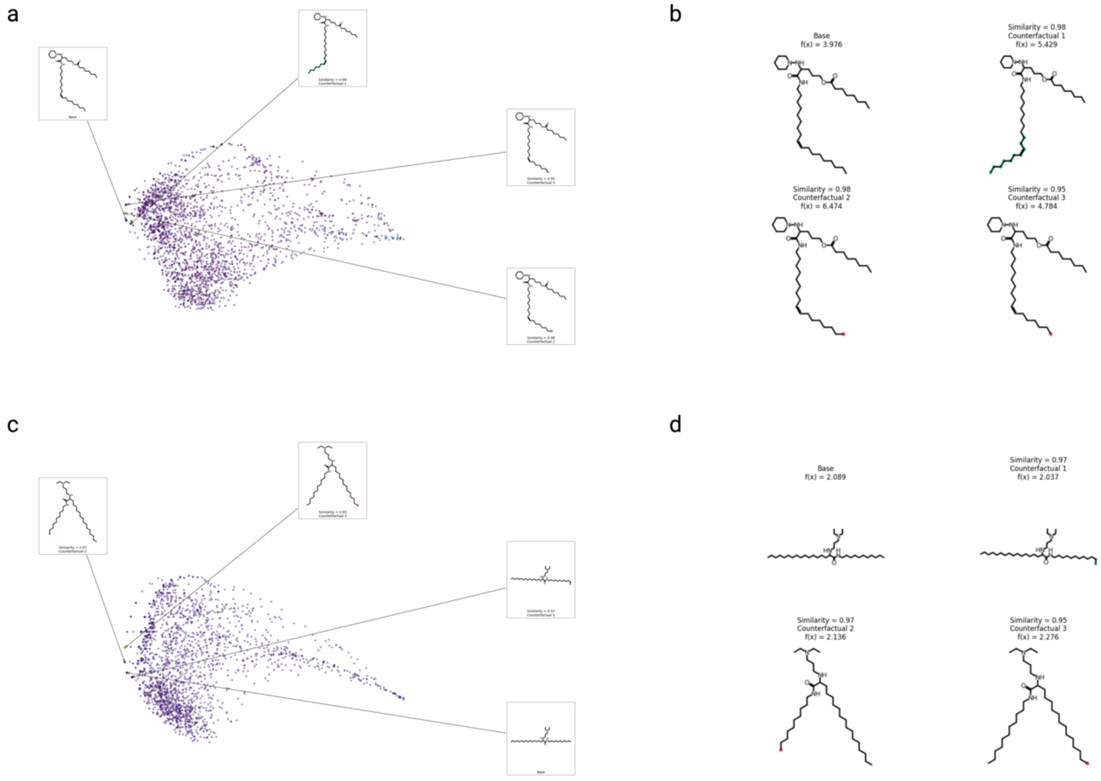


**Figure S22.** Illustration of Exmol molecular explanation using counterfactual generation, where the generated molecular counterfactuals are designed to retain as much similarity to the input lipid molecule as feasible. This approach can explore the specific regional changes that may lead to the escalation or decline of the predicted potencies. (a) The counterfactual space of H9. Each dot represents a counterfactual of H9. The base molecule is denoted as the farthest (left most one) from all other molecules. (b) A selection of four H9 counterfactuals are displayed, alongside an illustration of their similarities with the base molecule. (c) The counterfactual domain of R6 is depicted, with each point signifying a R6 counterfactual. The base molecule is marked as the one most distant from all other molecules (left most one). (d) A selection of four R6 counterfactuals are displayed, alongside an illustration of their similarities with the base molecule.

### Supplementary Notes

#### 1. Lipid synthesis and Characterization

##### Lipid Tails Synthesis for High-throughput Screening (1,200)

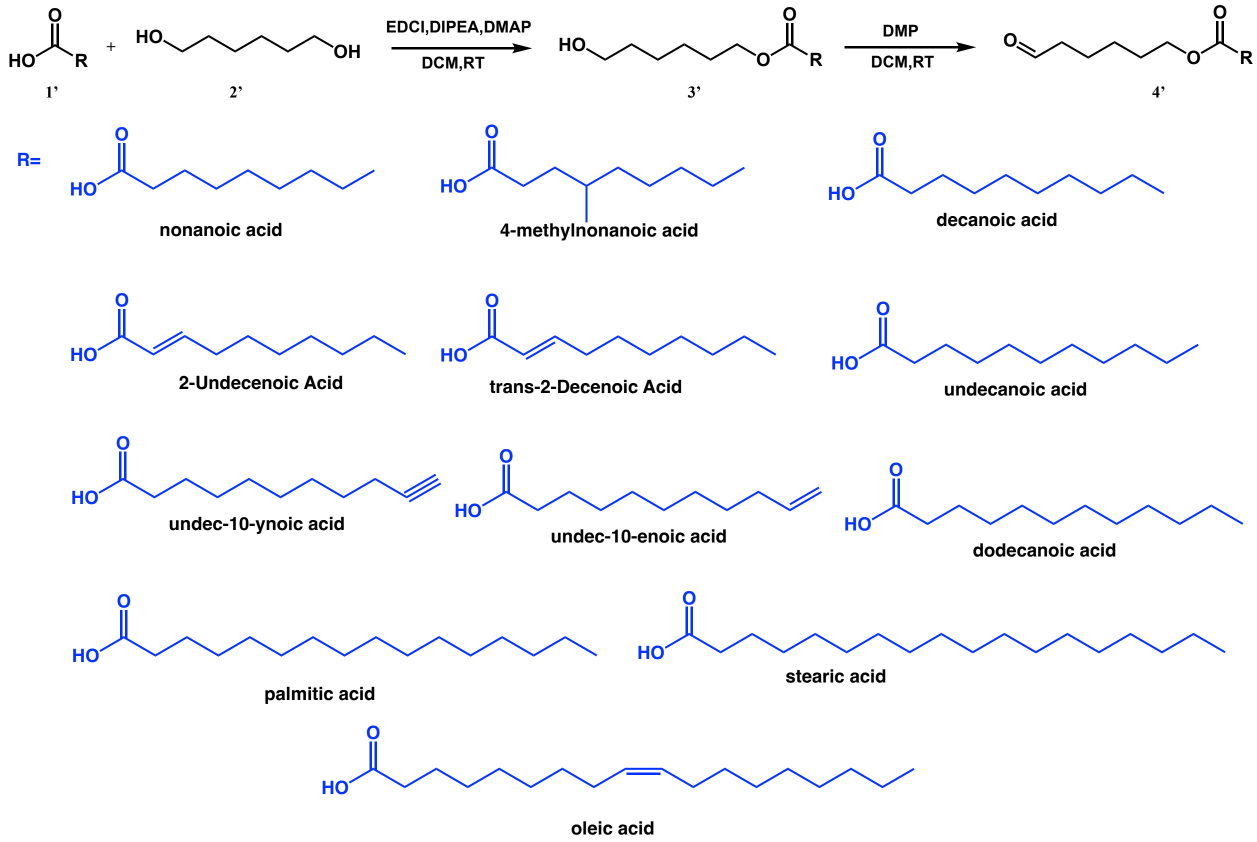


##### General Synthesis Route of Tail A

Acid (17.7 mmol), *N*-(3-dimethyl aminopropyl)- *N′*-ethyl carbodiimide hydrochloride (EDC HCl, 26.6 mmol), hexane-1,6-diol (88.5 mmol), 4-(dimethylamino) pyridine (DMAP, 8.9 mmol) and *N, N*-diisopropylethylamine (DIPEA, 35.4 mmol) were dissolved in dichloromethane (200 ml). The reaction was stirred at room temperature under nitrogen for 18 h, then washed with a saturated aqueous sodium bicarbonate solution. The organic layer was separated, washed with brine, dried over Na2SO4, filtered, and the filtrate was evaporated under a vacuum. The residue was purified by silica gel chromatography (0–50% ethyl acetate in hexanes) to give compound **3’**. **3’** (15.2 mmol) was dissolved in dichloromethane 300 ml followed by Dess-Martin Periodinane (9.6g, 22.8 mmol). The reaction was stirred under nitrogen at room temperature for 2 h. After confirmation of reaction completion by thin layer chromatography, sodium thiosulfate pentahydrate (50% w/v, 200 ml) was added to the reaction and left to stir for additional 15 mins. The organic layer was thereafter separated, washed with brine, dried over $Na_2SO_4$, filtered, and the filtrate was evaporated under a vacuum. The residue was purified by silica gel chromatography (0–50% ethyl acetate in hexanes) to give compound **4’**.

**<u>Synthesis of Tail 3’</u>**

<u>Synthesis of Tail 6-hydroxyhexyl nonanoate</u>

Follow the synthesis method above. $^1$H NMR (400 MHz, $CDCl_3$) δ 4.02 (t, *J* = 6.7 Hz, 2H), 3.59 (t, *J* = 6.6 Hz, 2H), 2.25 (t, *J* = 7.6 Hz, 2H), 1.67 – 1.47 (m, 6H), 1.35 (p, *J* = 4.0 Hz, 4H), 1.30 – 1.13 (m, 10H), 0.91 – 0.76 (m, 3H).

<u>Synthesis of Tail 6-hydroxyhexyl 4-methylnonanoate</u>

Follow the synthesis method above. $^1$H NMR (400 MHz, $CDCl_3$) δ 4.02 (dd, *J* = 7.2, 6.3 Hz, 2H), 3.59 (t, *J* = 6.6 Hz, 2H), 2.36 – 2.12 (m, 2H), 1.64 – 1.47 (m, 5H), 1.40 – 1.15 (m, 13H), 1.10 – 0.71 (m, 6H).

<u>Synthesis of Tail 6-hydroxyhexyl decanoate</u>

Follow the synthesis method above. $^1$H NMR (400 MHz, $CDCl_3$) δ 4.02 (t, *J* = 6.7 Hz, 2H), 3.58 (t, *J* = 6.6 Hz, 2H), 2.24 (t, *J* = 7.6 Hz, 2H), 1.73 – 1.46 (m, 6H), 1.39 – 1.31 (m, 4H), 1.26 – 1.17 (m, 11H), 0.85 – 0.80 (m, 3H).

<u>Synthesis of Tail 6-hydroxyhexyl undec-2-enoate</u>

Follow the synthesis method above. $^1$H NMR (400 MHz, $CDCl_3$) δ 6.92 (dt, *J* = 15.6, 7.0 Hz, 1H), 5.87 – 5.41 (m, 1H), 4.05 (dt, *J* = 19.2, 6.7 Hz, 2H), 3.58 (t, *J* = 6.6 Hz, 2H), 2.15 (qd, *J* = 7.1, 1.6 Hz, 2H), 1.72 – 1.46 (m, 4H), 1.45 – 1.31 (m, 6H), 1.25 – 1.21 (m, 6H), 0.94 – 0.69 (m, 3H).

<u>Synthesis of Tail 6-hydroxyhexyl (*E*)-dec-2-enoate</u>

Follow the synthesis method above. $^1$H NMR (400 MHz, $CDCl_3$) δ 6.91 (dt, *J* = 15.7, 7.0 Hz, 1H), 5.76 (dt, *J* = 15.6, 1.6 Hz, 1H), 4.04 (dt, *J* = 23.5, 6.7 Hz, 2H), 3.58 (t, *J* = 6.6 Hz, 2H), 2.27 – 2.06 (m, 2H), 1.76 – 1.46 (m, 5H), 1.45 – 1.30 (m, 6H), 1.24 – 1.19 (m, 6H), 0.89 – 0.76 (m, 3H).

<u>Synthesis of Tail 6-hydroxyhexyl undecanoate</u>

Follow the synthesis method above. $^1$H NMR (400 MHz, $CDCl_3$) δ 4.02 (t, *J* = 6.7 Hz, 2H), 3.59 (t, *J* = 6.6 Hz, 2H), 2.25 (t, *J* = 7.6 Hz, 2H), 1.70 – 1.45 (m, 6H), 1.41 – 1.31 (m, 4H), 1.30 – 1.13 (m, 14H), 0.93 – 0.71 (m, 3H).

<u>Synthesis of Tail 6-hydroxyhexyl undec-10-ynoate</u>

Follow the synthesis method above. $^1$H NMR (400 MHz, $CDCl_3$) δ 4.02 (t, *J* = 6.7 Hz, 2H), 3.59 (t, *J* = 6.6 Hz, 2H), 2.25 (t, *J* = 7.5 Hz, 2H), 2.13 (td, *J* = 7.1, 2.7 Hz, 2H), 1.90 (t, *J* = 2.6 Hz, 1H), 1.67 – 1.42 (m, 8H), 1.40 – 1.15 (m, 12H).

<u>Synthesis of Tail 6-hydroxyphenyl undec-10-enoate</u>

Follow the synthesis method above. $^1$H NMR (400 MHz, $CDCl_3$) δ 5.74 (ddt, *J* = 16.9, 10.2, 6.7 Hz, 1H), 5.04 – 4.77 (m, 2H), 4.00 (t, *J* = 6.7 Hz, 2H), 3.56 (t, *J* = 6.6 Hz, 2H), 2.23 (t, *J* = 7.5 Hz, 2H), 1.97 (dt, *J* = 8.0, 6.7 Hz, 2H), 1.70 – 1.43 (m, 6H), 1.41 – 0.97 (m, 14H).

<u>Synthesis of Tail 6-hydroxyhexyl dodecanoate</u>

Follow the synthesis method above. $^1$H NMR (400 MHz, $CDCl_3$) δ 4.02 (t, *J* = 6.7 Hz, 2H), 3.58 (t, *J* = 6.6 Hz, 2H), 2.25 (t, *J* = 7.6 Hz, 2H), 1.65 – 1.46 (m, 6H), 1.43 – 1.06 (m, 20H), 1.02 – 0.69 (m, 3H).

<u>Synthesis of Tail 6-hydroxyhexyl palmitate</u>

Follow the synthesis method above. $^1$H NMR (400 MHz, $CDCl_3$) δ 4.06 (t, *J* = 6.7 Hz, 2H), 3.64 (t, *J* = 6.5 Hz, 2H), 2.28 (t, *J* = 7.6 Hz, 2H), 1.60 (dddd, *J* = 14.5, 12.9, 6.8, 3.2 Hz, 6H), 1.38 (p, *J* = 3.3 Hz, 4H), 1.25 (s, 22H), 0.97 – 0.76 (m, 3H).

Synthesis of Tail 6-hydroxyhexyl stearate

Follow the synthesis method above. $^{1}$H NMR (400 MHz, $CDCl_3$) δ 4.04 (t, *J* = 6.7 Hz, 2H), 3.62 (t, *J* = 6.6 Hz, 2H), 2.27 (t, *J* = 7.6 Hz, 2H), 1.59 (dddd, *J* = 15.1, 9.8, 6.7, 3.7 Hz, 7H), 1.37 (p, *J* = 3.8, 3.3 Hz, 4H), 1.23 (s, 28H), 0.90 – 0.77 (m, 3H).

Synthesis of Tail 6-hydroxyhexyl oleate

Follow the synthesis method above. $^{1}$H NMR (400 MHz, $CDCl_3$) δ 5.46 – 5.21 (m, 2H), 4.04 (t, *J* = 6.7 Hz, 2H), 3.62 (t, *J* = 6.6 Hz, 2H), 2.27 (t, *J* = 7.6 Hz, 2H), 2.06 – 1.85 (m, 4H), 1.63 – 1.53 (m, 6H), 1.41 – 1.34 (m, 4H), 1.33 – 1.16 (m, 20H), 0.97 – 0.79 (m, 3H).

**Synthesis of Tail 4'**

Synthesis of Tail 6-oxohexyl nonanoate

Follow the synthesis method above. $^{1}$H NMR (400 MHz, $CDCl_3$) δ 9.76 (t, *J* = 1.7 Hz, 1H), 4.05 (t, *J* = 6.6 Hz, 2H), 2.57 – 2.07 (m, 4H), 1.75 – 1.49 (m, 6H), 1.46 – 1.33 (m, 2H), 1.32 – 1.18 (m, 11H), 0.91 – 0.80 (m, 3H).

Synthesis of Tail 6-oxohexyl 4-methylnonanoate

Follow the synthesis method above. $^{1}$H NMR (400 MHz, $CDCl_3$) δ 9.76 (t, *J* = 1.7 Hz, 1H), 4.05 (t, *J* = 6.6 Hz, 2H), 2.48 – 2.26 (m, 4H), 1.69 – 1.54 (m, 6H), 1.40 (tdd, *J* = 8.2, 6.5, 4.5 Hz, 4H), 1.29 – 1.21 (m, 8H), 0.86 (td, *J* = 6.6, 1.3 Hz, 6H).

Synthesis of Tail 6-oxohexyl decanoate

Follow the synthesis method above. $^{1}$H NMR (400 MHz, $CDCl_3$) δ 9.74 (t, *J* = 1.7 Hz, 1H), 4.04 (t, *J* = 6.6 Hz, 2H), 2.55 – 2.14 (m, 4H), 1.62 (ddq, *J* = 14.3, 9.9, 7.3 Hz, 6H), 1.43 – 1.13 (m, 14H), 0.92 – 0.78 (m, 3H).

Synthesis of Tail 6-oxohexyl undec-2-enoate

Follow the synthesis method above. $^{1}$H NMR (400 MHz, $CDCl_3$) δ 9.76 (t, *J* = 1.7 Hz, 1H), 6.95 (dt, *J* = 15.6, 7.0 Hz, 1H), 5.79 (dt, *J* = 15.6, 1.6 Hz, 1H), 4.12 (t, *J* = 6.6 Hz, 2H), 2.48 – 2.25 (m, 3H), 2.18 (qd, *J* = 7.1, 1.6 Hz, 2H), 1.68 – 1.62 (m, 4H), 1.45 – 1.35 (m, 5H), 1.26 – 1.23 (m, 6H), 0.86 (d, *J* = 7.1 Hz, 3H).

Synthesis of Tail 6-oxohexyl (E)-dec-2-enoate

Follow the synthesis method above. $^{1}$H NMR (400 MHz, $CDCl_3$) δ 9.82 – 9.66 (m, 1H), 6.93 (dt, *J* = 15.6, 6.9 Hz, 1H), 5.77 (dt, *J* = 15.6, 1.8 Hz, 1H), 4.09 (t, *J* = 6.6 Hz, 2H), 2.43 (td, *J* = 7.3, 1.7 Hz, 2H), 2.16 (qd, *J* = 7.3, 1.6 Hz, 2H), 1.71 – 1.61 (m, 4H), 1.46 – 1.35 (m, 4H), 1.26 – 1.18 (m, 6H), 0.84 (t, *J* = 6.7 Hz, 3H).

Synthesis of Tail 6-oxohexyl undecanoate

Follow the synthesis method above. $^{1}$H NMR (400 MHz, $CDCl_3$) δ 9.74 (t, *J* = 1.7 Hz, 1H), 4.04 (t, *J* = 6.6 Hz, 2H), 2.43 (td, *J* = 7.3, 1.7 Hz, 2H), 2.26 (t, *J* = 7.6 Hz, 2H), 1.61 (ddt, *J* = 16.2, 9.8, 7.3 Hz, 6H), 1.42 – 1.34 (m, 2H), 1.24 (d, *J* = 9.2 Hz, 14H), 0.87 – 0.82 (m, 3H).

Synthesis of Tail 6-oxohexyl undec-10-ynoate

Follow the synthesis method above. $^{1}$H NMR (400 MHz, $CDCl_3$) δ 4.02 (t, *J* = 6.6 Hz, 2H), 2.41 (td, *J* = 7.3, 1.7 Hz, 1H), 2.32 (t, *J* = 7.5 Hz, 1H), 2.24 (t, *J* = 7.5 Hz, 2H), 2.13 (td, *J* = 7.1, 2.7 Hz,

2H), 1.90 (t, $J$ = 2.7 Hz, 1H), 1.65 – 1.55 (m, 6H), 1.52 – 1.43 (m, 2H), 1.39 – 1.31 (m, 4H), 1.27 – 1.17 (m, 6H).

<u>Synthesis of Tail 6-oxohexyl undec-10-enoate</u>

Follow the synthesis method above. $^1$H NMR (400 MHz, $CDCl_3$) δ 9.76 (t, $J$ = 1.7 Hz, 1H), 5.79 (ddt, $J$ = 16.9, 10.2, 6.7 Hz, 1H), 5.07 – 4.79 (m, 2H), 4.06 (t, $J$ = 6.6 Hz, 2H), 2.45 (td, $J$ = 7.3, 1.7 Hz, 2H), 2.28 (t, $J$ = 7.5 Hz, 2H), 2.08 – 1.95 (m, 2H), 1.71 – 1.55 (m, 6H), 1.45 – 1.37 (m, 2H), 1.36 – 1.12 (m, 9H).

<u>Synthesis of Tail 6-oxohexyl dodecanoate</u>

Follow the synthesis method above. $^1$H NMR (400 MHz, $CDCl_3$) δ 9.73 (t, $J$ = 1.8 Hz, 1H), 4.03 (t, $J$ = 6.6 Hz, 2H), 2.42 (td, $J$ = 7.3, 1.7 Hz, 2H), 2.25 (t, $J$ = 7.5 Hz, 2H), 1.62 (dd, $J$ = 15.3, 7.4 Hz, 6H), 1.37 (ddt, $J$ = 9.3, 6.7, 3.4 Hz, 2H), 1.22 (s, 16H), 0.83 (d, $J$ = 7.0 Hz, 3H).

<u>Synthesis of Tail 6-oxohexyl palmitate</u>

Follow the synthesis method above. $^1$H NMR (400 MHz, $CDCl_3$) δ 9.76 (t, $J$ = 1.6 Hz, 1H), 4.05 (t, $J$ = 6.6 Hz, 2H), 2.49 – 2.34 (m, 2H), 2.28 (t, $J$ = 7.5 Hz, 2H), 1.72 – 1.56 (m, 6H), 1.39 (tt, $J$ = 9.8, 6.3 Hz, 2H), 1.24 (s, 22H), 0.87 (t, $J$ = 6.7 Hz, 3H).

<u>Synthesis of Tail 6-oxohexyl stearate</u>

Follow the synthesis method above. $^1$H NMR (400 MHz, $CDCl_3$) δ 9.76 (t, $J$ = 1.7 Hz, 1H), 4.05 (t, $J$ = 6.6 Hz, 2H), 2.44 (td, $J$ = 7.3, 1.7 Hz, 2H), 2.28 (t, $J$ = 7.6 Hz, 2H), 1.72 – 1.53 (m, 6H), 1.48 – 1.06 (m, 32H), 0.91 – 0.78 (m, 3H).

<u>Synthesis of Tail 6-oxohexyl oleate</u>

Follow the synthesis method above. $^1$H NMR (400 MHz, $CDCl_3$) δ 9.75 (t, $J$ = 1.7 Hz, 1H), 5.37 – 5.27 (m, 2H), 4.04 (t, $J$ = 6.6 Hz, 2H), 2.43 (td, $J$ = 7.3, 1.7 Hz, 2H), 2.27 (t, $J$ = 7.6 Hz, 2H), 1.99 (q, $J$ = 6.7 Hz, 3H), 1.71 – 1.54 (m, 6H), 1.43 – 1.34 (m, 2H), 1.32 – 1.19 (m, 20H), 0.87 – 0.81 (m, 3H).

**<u>General Synthesis Route of Tail B</u>**

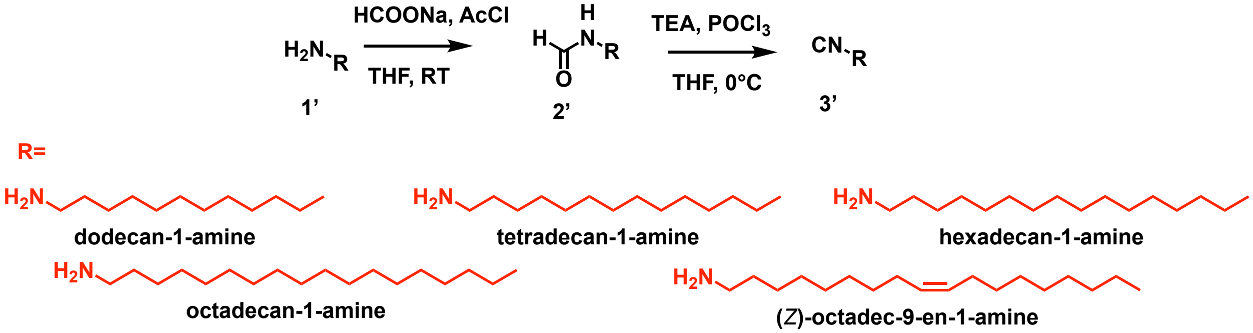


Tetrahydrofuran (82.75 ml) and solid sodium formate (2.2511 g) were added into a 250 mL round-bottomed flask and stirred to form a suspension. A DCM (34.755 mL) solution containing acetyl chloride (1.0 M) was taken in a constant pressure titration funnel and added dropwise to the suspension prepared above. The mixture was stirred at room temperature for 12 h after the dropwise addition to obtain reaction solution 1. 1' was dissolved in THF and added dropwise to reaction solution 1. The mixture was stirred at room temperature for 4 h after the dropwise addition to obtain reaction solution 2. Reaction solution 2 was diluted with 125 ml of deionized water, and the aqueous solution was extracted twice with 240 ml of ethyl acetate, each with an amount of 120 ml. The resulting extract was dried with anhydrous magnesium sulphate for 0.5 h and spin dried to obtain the crude 2'.

The crude 2' product and THF (50 mL) solution were added to a round-bottomed flask (100 mL) and placed in an ice bath at 0°C. The mixture was stirred for 10 min and TEA was added to the solution. The mixture was stirred in the ice bath for 15 min to obtain the reaction solution 1. THF (12.9 mL) and phosphorus oxychloride (3.519 g) were taken to prepare solution 1. Under ice bath conditions, solution 1 was added slowly dropwise to reaction solution 1 with a constant pressure-dropping funnel. The reaction was stirred for 1.5 h. 65 mL of a saturated sodium bicarbonate solution was added to dilute and wash the mixture, and the washed solution was extracted twice with 130 mL of ether, 65 mL each time. The extracts were combined, and anhydrous magnesium sulphate was added to the organic phase, stirred and dried for 0.5 h. The solution was removed by rotary evaporation. The crude product was purified by column using a Hexane/EA system.

Synthesis of Tail 1-isocyanododecane

Follow the synthesis method above. $^1$H NMR (400 MHz, $CDCl_3$) δ 3.49 – 3.22 (m, 2H), 1.72 – 1.57 (m, 2H), 1.50 – 1.34 (m, 2H), 1.26 (d, *J* = 10.0 Hz, 16H), 0.89 – 0.84 (m, 3H).

Synthesis of Tail 1-isocyanotetradecane

Follow the synthesis method above. $^1$H NMR (400 MHz, $CDCl_3$) δ 3.47 – 3.24 (m, 2H), 1.66 (ddt, *J* = 10.8, 8.4, 4.2 Hz, 2H), 1.46 – 1.37 (m, 2H), 1.27 (d, *J* = 11.2 Hz, 20H), 0.94 – 0.74 (m, 3H).

Synthesis of Tail 1-isocyanohexadecane

Follow the synthesis method above. $^1$H NMR (400 MHz, $CDCl_3$) δ 3.45 – 3.26 (m, 2H), 1.66 (ddtt, *J* = 11.4, 6.7, 4.6, 2.3 Hz, 2H), 1.46 – 1.38 (m, 2H), 1.26 (d, *J* = 11.1 Hz, 24H), 0.87 (t, *J* = 6.8 Hz, 3H).

Synthesis of Tail 1-isocyanooctadecane

Follow the synthesis method above. $^1$H NMR (400 MHz, $CDCl_3$) δ 3.48 – 3.23 (m, 2H), 1.73 – 1.61 (m, 2H), 1.46 – 1.37 (m, 2H), 1.25 (s, 28H), 0.91 – 0.84 (m, 3H).

Synthesis of Tail (Z)-1-isocyanooctadec-9-ene

Follow the synthesis method above. $^1$H NMR (400 MHz, $CDCl_3$) δ 5.41 – 5.21 (m, 2H), 3.45 – 3.29 (m, 2H), 1.99 (dq, *J* = 14.2, 6.0 Hz, 3H), 1.66 (ddd, *J* = 8.4, 5.6, 3.4 Hz, 2H), 1.48 – 1.35 (m, 2H), 1.28 (dt, *J* = 17.3, 6.2 Hz, 20H), 0.92 – 0.84 (m, 3H).

**Tail Synthesis for ML-predicted structure**

Synthesis of olealdehyde

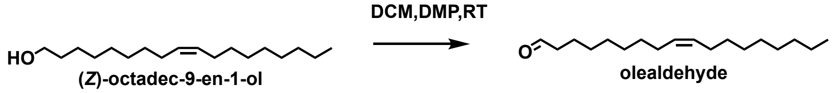


Olealdehyde was made using the same procedure as the synthesis method above. $^1$H NMR (400 MHz, $CDCl_3$) δ 9.75 (t, *J* = 1.9 Hz, 1H), 5.34 (qd, *J* = 3.8, 1.7 Hz, 2H), 2.41 (td, *J* = 7.4, 1.9 Hz, 2H), 2.00 (q, *J* = 6.1 Hz, 4H), 1.62 (p, *J* = 7.3 Hz, 2H), 1.34 – 1.22 (m, 20H), 0.91 – 0.83 (m, 3H).

Synthesis of palmitaldehyde

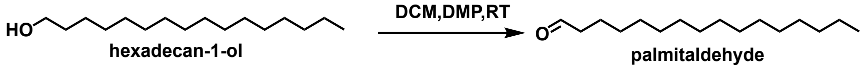


Palmitaldehyde was made using the same procedure as the synthesis method above. $^1$H NMR (400 MHz, $CDCl_3$) δ 9.76 (s, 1H), 2.41 (td, *J* = 7.4, 1.9 Hz, 2H), 1.62 (dd, *J* = 9.3, 5.2 Hz, 2H), 1.25 (s, 23H), 0.90 – 0.84 (m, 3H).

Synthesis of 4-oxobutyl 4-methylnonanoate

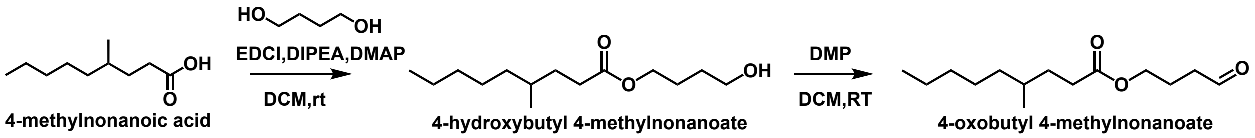


4-hydroxybutyl 4-methylnonanoate was made using the same procedure as the synthesis method above. $^1$H NMR (400 MHz, $CDCl_3$) δ 4.09 – 4.04 (m, 2H), 3.64 (t, *J* = 6.3 Hz, 2H), 2.27 (td, *J* = 9.3, 6.1 Hz, 2H), 1.70 – 1.57 (m, 4H), 1.45 – 1.15 (m, 10H), 0.90 – 0.77 (m, 6H). 4-oxobutyl 4-methylnonanoate was made using the same procedure as the synthesis method above. $^1$H NMR (400 MHz, $CDCl_3$) δ 9.78 (s, 1H), 4.08 (t, *J* = 6.3 Hz, 2H), 2.53 (td, *J* = 7.2, 1.3 Hz, 2H), 2.28 (td, *J* = 9.4, 6.1 Hz, 2H), 2.01 – 1.90 (m, 2H), 1.66 – 1.57 (m, 1H), 1.44 – 1.19 (m, 10H), 0.88 – 0.83 (m, 6H).

Synthesis of 4-oxobutyl (E)-dec-2-enoate

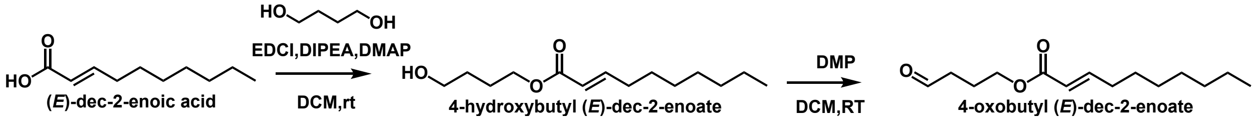


4-hydroxybutyl (*E*)-dec-2-enoate was made using the same procedure as the synthesis method above. $^1$H NMR (400 MHz, $CDCl_3$) δ 6.89 (dt, *J* = 15.6, 6.9 Hz, 1H), 5.73 (dt, *J* = 15.7, 1.6 Hz, 1H), 4.08 (t, *J* = 6.5 Hz, 2H), 3.58 (t, *J* = 6.4 Hz, 2H), 2.12 (qd, *J* = 7.1, 1.6 Hz, 2H), 1.71 – 1.54 (m, 5H), 1.41 – 1.34 (m, 2H), 1.24 – 1.19 (m, 8H), 0.80 (d, *J* = 7.1 Hz, 3H).
4-oxobutyl (*E*)-dec-2-enoate was made using the same procedure as the synthesis method above. $^1$H NMR (400 MHz, $CDCl_3$) δ 9.78 (t, *J* = 1.3 Hz, 1H), 6.95 (dt, *J* = 15.6, 6.9 Hz, 1H), 5.78 (d, *J* = 15.7 Hz, 1H), 4.15 (t, *J* = 6.3 Hz, 2H), 2.55 (td, *J* = 7.2, 1.3 Hz, 2H), 2.18 (qd, *J* = 7.1, 1.6 Hz, 2H), 1.99 (ddd, *J* = 13.6, 7.3, 6.3 Hz, 2H), 1.46 – 1.38 (m, 2H), 1.29 – 1.23 (m, 8H), 0.88 – 0.84 (m, 3H).

Synthesis of 4-oxobutyl octanoate

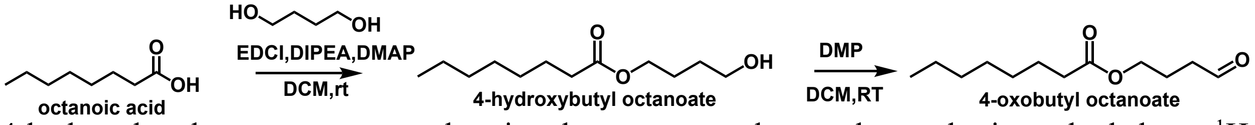


4-hydroxybutyl octanoate was made using the same procedure as the synthesis method above. $^1$H NMR (400 MHz, $CDCl_3$) δ 4.06 (q, *J* = 7.1 Hz, 2H), 3.67 (p, *J* = 6.7 Hz, 1H), 3.08 (q, *J* = 7.4 Hz, 1H), 2.23 – 2.19 (m, 3H), 1.31 (d, *J* = 6.8 Hz, 7H), 1.22 – 1.20 (m, 9H), 0.82 (d, *J* = 4.0 Hz, 6H). 4-oxobutyl octanoate was made using the same procedure as the synthesis method above. $^1$H NMR (400 MHz, $CDCl_3$) δ 9.77 (t, *J* = 1.3 Hz, 1H), 4.08 (t, *J* = 6.3 Hz, 2H), 2.53 (td, *J* = 7.2, 1.3 Hz,

2H), 2.30 – 2.22 (m, 2H), 1.96 (ddd, $J$ = 13.5, 7.2, 6.3 Hz, 2H), 1.67 – 1.53 (m, 2H), 1.31 – 1.21 (m, 9H), 0.88 – 0.83 (m, 3H).

Synthesis of 4-oxobutyl oleate

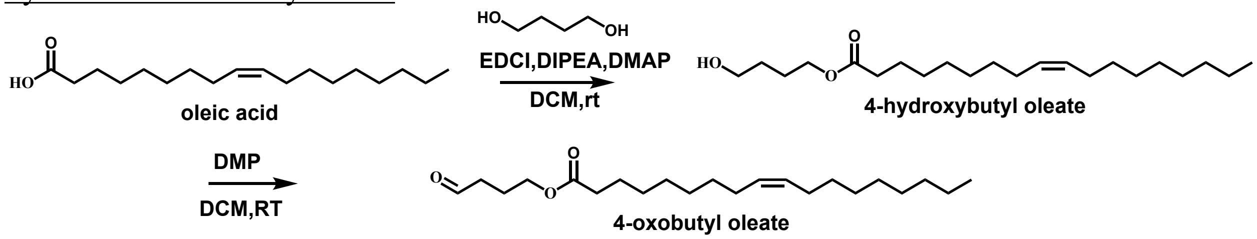


4-hydroxybutyl oleate was made using the same procedure as the synthesis method above. $^1$H NMR (400 MHz, $CDCl_3$) δ 5.42 – 5.25 (m, 2H), 4.12 – 4.09 (m, 2H), 3.68 (t, $J$ = 6.3 Hz, 2H), 2.29 (t, $J$ = 7.6 Hz, 2H), 2.00 (q, $J$ = 6.5 Hz, 4H), 1.76 – 1.60 (m, 6H), 1.31 – 1.24 (m, 20H), 0.87 (t, $J$ = 6.7 Hz, 3H). 4-oxobutyl oleate was made using the same procedure as the synthesis method above. $^1$H NMR (400 MHz, $CDCl_3$) δ 9.70 (t, $J$ = 1.3 Hz, 1H), 5.45 – 5.15 (m, 2H), 4.14 (m, 2H), 2.52 – 2.18 (m, 4H), 2.12 – 1.78 (m, 6H), 1.70 – 1.50 (m, 2H), 1.41 – 1.11 (m, 20H), 1.02 – 0.78 (m, 3H).

Synthesis of 1-isocyanoundecane

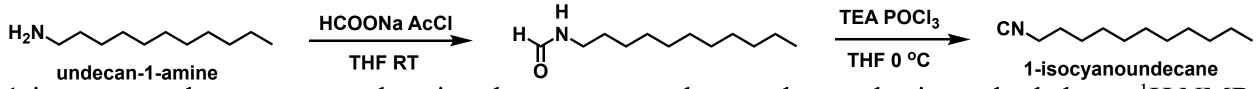


1-isocyanoundecane was made using the same procedure as the synthesis method above. $^1$H NMR (400 MHz, $CDCl_3$) δ 3.42 – 3.27 (m, 2H), 1.63 – 1.52 (m, 2H), 1.31 – 1.24 (m, 16H), 0.88 – 0.86 (m, 3H).

Synthesis of 1-isocyanoheptadecane

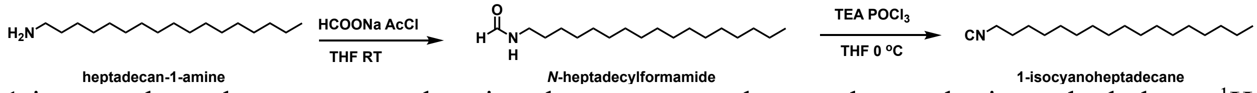


1-isocyanoheptadecane was made using the same procedure as the synthesis method above. $^1$H NMR (400 MHz, $CDCl_3$) δ 3.44 – 3.29 (m, 2H), 1.74 – 1.37 (m, 4H), 1.25 (d, $J$ = 1.5 Hz, 15H), 0.87 (td, $J$ = 6.9, 1.8 Hz, 3H).

**The mechanism of the Ugi-3CR reaction is mediated by isocyanide and its Markush structure.**

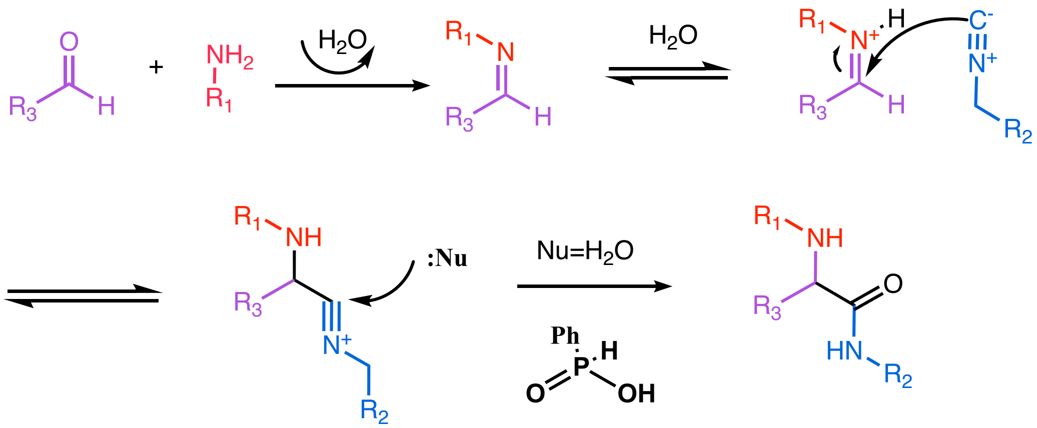


**Top-performing lipid synthesis**

Synthesis Method of Lipid **H9** (*Z*)-5-(octadec-9-en-1-ylamino)-5-oxo-4-(piperidin-1-ylamino)pentyl octanoate.

The synthesis of **H9** through 3CR-Ugi reaction. A mixture of piperidin-1-amine (25 mg, 249.59 µmol), 4-oxobutyl octanoate (53.49 mg, 249.59 µmol) and (Z)-1-isocyanooctadec-9-ene (69.26 mg, 249.59 µmol) in anhydrous solvent DCM (240 µL) and MeOH (160 µL) with the catalyst phosphoric acid (7.09 mg, 49.92 µmol) and was stirred in capped glass vials at room temperature overnight. The mixture was purified by flash column chromatography (0.1% ammonia hydroxide with 0-10% MeOH in DCM) to obtain compound **H9** as colorless oil. (35.0 mg, 59.13 µmol, 23.69%). $^1$H NMR (400 MHz, $CDCl_3$) δ 7.03 (t, *J* = 5.8 Hz, 1H), 5.46 – 5.31 (m, 2H), 4.05 (t, *J* = 6.1 Hz, 2H), 3.40 – 3.26 (m, 2H), 3.23 – 3.11 (m, 1H), 2.69 (s, 2H), 2.41 (s, 2H), 2.28 (t, *J* = 7.6 Hz, 2H), 1.99 (dd, *J* = 14.1, 7.7 Hz, 4H), 1.74 – 1.46 (m, 14H), 1.33 – 1.21 (m, 31H), 0.91 – 0.83 (m, 6H). Chemical Formula: $C_{36}H_{69}N_3O_3$ and MS calculated: m/z 591.53, observed (ESI ms) m/z 592.53.

Synthesis Method of Lipid **R6** 2-((3-(diethylamino)propyl) amino) -N-undecylheptadecanamide.

The synthesis of **R6** through 3CR-Ugi reaction. A mixture of *N*, *N*-diethylpropane-1,3-diamine (25 mg, 191.96 µmol), palmitaldehyde (46.15 mg, 191.96 µmol) and 1-isocyanoundecane (34.81 mg, 191.96 µmol) in anhydrous solvent DCM (240 µL) and MeOH (160 µL) with the catalyst phosphoric acid (5.46 mg, 39.39 µmol) and was stirred in capped glass vials at room temperature overnight. The mixture was purified by flash column chromatography (0.1% ammonia hydroxide with 0-10% MeOH in DCM) to obtain compound **R6** as colorless oil. (39.8 mg, 72.1 µmol, 37.56%). $^1$H NMR (400 MHz, $CDCl_3$) δ 3.27 – 3.18 (m, 2H), 2.99 (dd, *J* = 7.7, 4.7 Hz, 1H), 2.67 – 2.40 (m, 8H), 1.73 – 1.43 (m, 6H), 1.33 – 1.19 (m, 44H), 1.03 (td, *J* = 7.1, 3.1 Hz, 7H), 0.91 – 0.84 (m, 6H). Chemical Formula: $C_{35}H_{73}N_3O$ and MS calculated: m/z 551.58, observed (ESI ms) m/z 552.73.

**NMR and Mass Spectrometry Assignments for Top Performing Lipids**

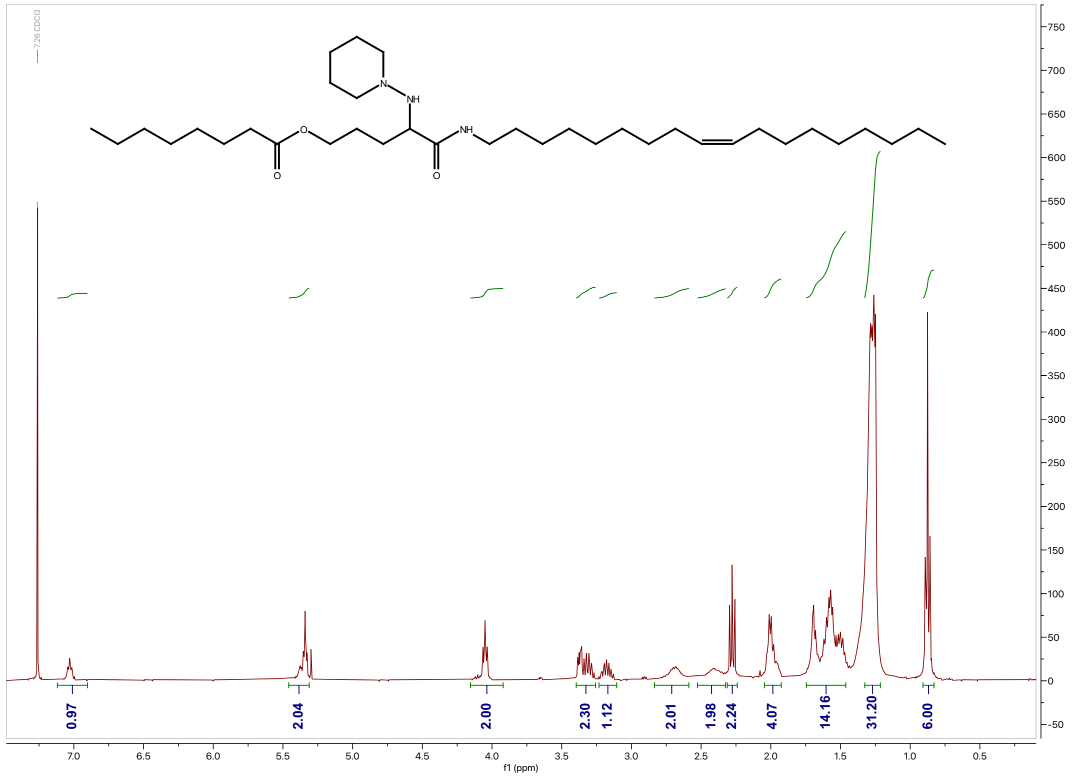


[1]H NMR of **H9**.

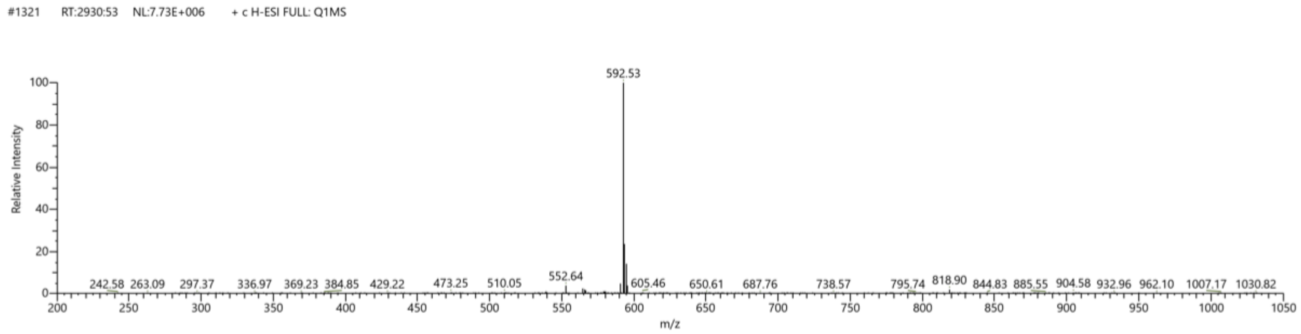


Mass Spectrum of **H9**.

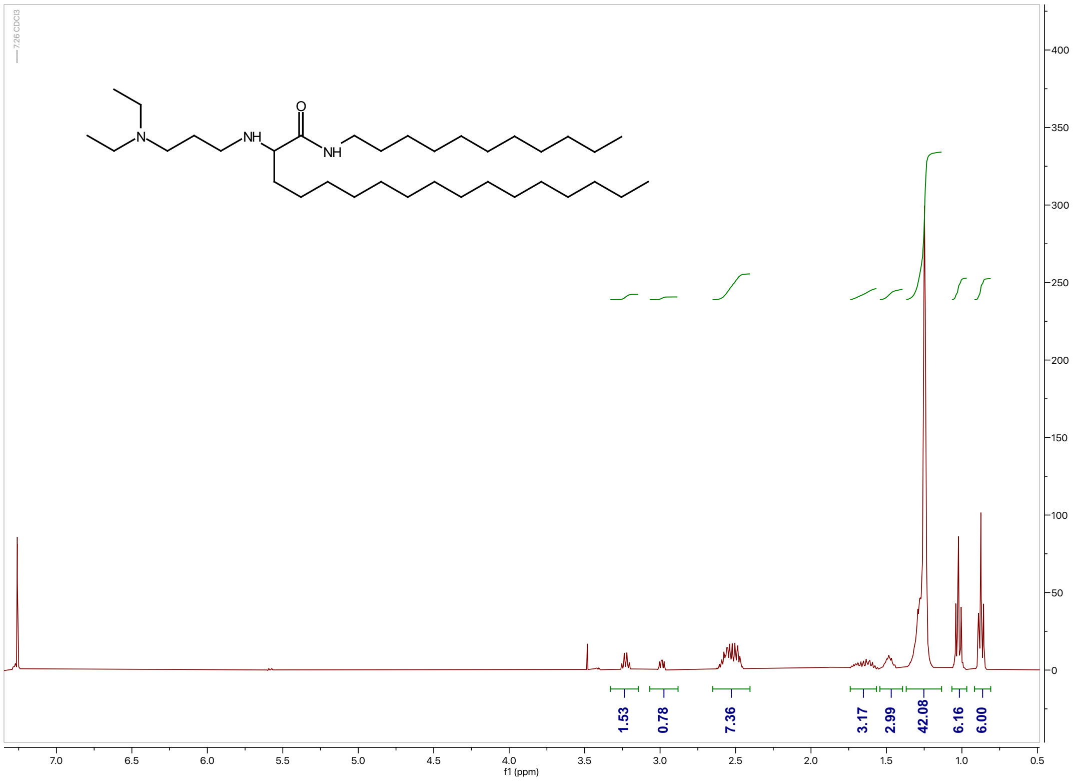


[1]H NMR of **R6**.

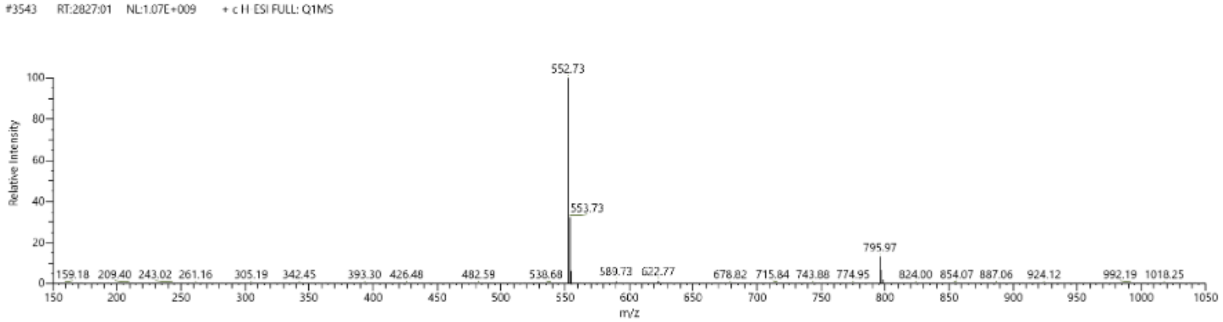


Mass Spectrum of **R6**.

#### Supplementary Tables

**Table S1. Formulation and characterization for LNP formulations with the top-performing ionizable lipids (H9 and R6)**

| | Factor | | | | Response |
|---|---|---|---|---|---|
| Nmuber | Lipid/mRNA weight ratio | Ionizable Lipid (mol%) | DOPE (mol%) | PEG (mol%) | In Vitro Hela ($Log_2$) |
| 1 | 7.5 | 60 | 20 | 1.5 | 9.246 |
| 2 | 7.5 | 60 | 15 | 0.5 | 11.086 |
| 3 | 15 | 30 | 10 | 1.5 | 4.466 |
| 4 | 7.5 | 30 | 20 | 2.5 | 8.412 |
| 5 | 7.5 | 30 | 10 | 2.5 | 2.616 |
| 6 | 7.5 | 60 | 10 | 2.5 | -0.071 |
| 7 | 15 | 60 | 10 | 2.5 | 8.257 |
| *8 | 15 | 60 | 20 | 0.5 | 12.244 |
| 9 | 15 | 60 | 10 | 0.5 | 8.169 |
| 10 | 11.25 | 60 | 20 | 2.5 | 9.016 |
| 11 | 15 | 45 | 20 | 2.5 | 8.923 |
| 12 | 7.5 | 45 | 10 | 0.5 | 6.256 |
| 13 | 15 | 30 | 20 | 0.5 | 9.252 |
| 14 | 11.25 | 45 | 15 | 1.5 | 8.382 |
| 15 | 15 | 30 | 15 | 2.5 | 4.237 |
| 16 | 7.5 | 30 | 20 | 0.5 | 8.024 |
| 17 | 11.25 | 30 | 10 | 0.5 | 6.939 |

**Note: The LNP formulation with the best performance for H9 was *8.**

| Nmuber | Factor | | | | Response |
|---|---|---|---|---|---|
| | Lipid/mRNA weight ratio | Ionizable Lipid (mol%) | DOTAP (mol%) | PEG (mol%) | In Vitro RAW 264.7 ($Log_2$) |
| 1 | 7.5 | 60 | 20 | 1.5 | 6.518 |
| #2 | 7.5 | 60 | 15 | 0.5 | 6.680 |
| 3 | 15 | 30 | 10 | 1.5 | 4.327 |
| 4 | 7.5 | 30 | 20 | 2.5 | 4.414 |
| 5 | 7.5 | 30 | 10 | 2.5 | 2.705 |
| 6 | 7.5 | 60 | 10 | 2.5 | 2.825 |
| 7 | 15 | 60 | 10 | 2.5 | 3.102 |
| 8 | 15 | 60 | 20 | 0.5 | 5.172 |
| 9 | 15 | 60 | 10 | 0.5 | 5.574 |
| 10 | 11.25 | 60 | 20 | 2.5 | 5.115 |
| 11 | 15 | 45 | 20 | 2.5 | 4.621 |
| 12 | 7.5 | 45 | 10 | 0.5 | 6.123 |
| 13 | 15 | 30 | 20 | 0.5 | 5.604 |
| 14 | 11.25 | 45 | 15 | 1.5 | 3.333 |
| 15 | 15 | 30 | 15 | 2.5 | 3.628 |
| 16 | 7.5 | 30 | 20 | 0.5 | 6.443 |
| 17 | 11.25 | 30 | 10 | 0.5 | 5.150 |

**Note: Formulation #2 was identified as the top performing LNP formulation for R6.**
